# HDL Nanodiscs Loaded with Liver X Receptor Agonist Decreases Tumor Burden and Mediates Long-term Survival in Mouse Glioma Model

**DOI:** 10.1101/2025.04.01.646644

**Authors:** Troy A. Halseth, Anzar A. Mujeeb, Lisha Liu, Kaushik Banerjee, Nigel Lang, Todd Hollon, Minzhi Yu, Mark Vander Roest, Ling Mei, Hongliang He, Maya Sheth, Maria G. Castro, Anna Schwendeman

## Abstract

Glioblastoma multiforme (GBM) is highly aggressive primary brain tumor with a 5-year survival rate of 7%. Previous studies have shown that GBM tumors have a reduced capacity to produce cholesterol and instead depend on the uptake of cholesterol produced by astrocytes. To target cholesterol metabolism to induce cancer cell death, synthetic high-density lipoprotein (sHDL) nanodiscs delivering Liver-X-Receptor (LXR) agonists and CpG oligonucleotides for targeting GBM were investigated. LXR agonists synergize with sHDL nanodiscs by increasing the expression of the ABCA1 cholesterol CpG oligonucleotides are established adjuvants used in cancer immunotherapy that work through the toll-like receptor 9 pathway. In the present study, treatment with GW-CpG-sHDL nanodiscs increased the expression of cholesterol efflux transporters on murine GL261 cells leading to enhanced cholesterol removal. Experiments in GL261-tumor-bearing mice revealed combining GW-CpG-sHDL nanodiscs with radiation (IR) therapy significantly increases median survival compared to GW-CpG-sHDL or IR alone. Furthermore, 66% of long-term survivors from the GW-CpG-sHDL +IR treatment group showed no tumor tissue when rechallenged.

## 1. Introduction

The current standard of care for GBM includes surgical resection, radiation therapy, and chemotherapy with temozolomide.^[1, 2]^ Surgical resection is limited in utility due to the highly invasive nature of GBM tumors. Furthermore, treatment with temozolomide only increases survival by 2-3 months beyond radiation and surgery and is also limited by drug resistance in tumors.^[1, 2]^ Because of this, it is critical to investigate alternative treatment strategies for GBM.

Recent areas of research in GBM treatments include immune modulation to overcome the suppressive tumor microenvironment, as well as treatments targeting cholesterol metabolism.^[3–8]^ Previous groups have demonstrated that administering nanoparticles loaded with CpG, a toll-like receptor (TLR) agonist, resulted in potent immune stimulation and significantly increased survival in glioma-bearing mice.^[9]^ Other researchers have investigated cholesterol as a therapeutic target in GBM in addition to direct immune cell stimulation. Cholesterol accumulation in the tumor microenvironment has been shown to induce CD8+ T-cell exhaustion by upregulating the expression of inhibitory receptors such as PD-1 and 2B4.^[10]^ Importantly, previous studies have shown that treatment of T-cells with B-cyclodextrin (which binds and removes cholesterol from cells) can reverse T-cell exhaustion. Targeting cholesterol is particularly attractive in GBM because of the alterations in the expression of several genes related to cholesterol synthesis and transport. It has been shown that GBM cells have reduced expression of *de novo* cholesterol synthesis genes but express high levels of the low-density lipoprotein receptor (LDLR), which is important for the uptake of cholesterol synthesized by astrocytes.^[4, 5, 8]^ This increased uptake of cholesterol enables the tumor cells to proliferate more freely. Because cholesterol is the primary energy source for tumor cells and is essential for tumor cell survival, strategies aimed at lowering cholesterol levels have been investigated for targeting GBM.^[11–15]^ Villa et al. have shown that treating GBM mice with the liver X receptor (LXR) agonist, LXR-623, reduced tumor growth and increased the survival of glioma-bearing mice. LXR agonists bind to nuclear receptors, increase the expression of cholesterol efflux transporters ABCA1 and ABCG1, and trigger the degradation of LDLR through upregulation of IDOL E3 ligase.^[13, 14, 16–18]^ These mechanisms all work in concert to lower intracellular cholesterol levels and deprive the tumor cells of cholesterol and trigger tumor cell death in GBM.

In the present study, we sought to combine the strategies of immune stimulation and reduction of cholesterol levels by co-delivery of CpG and the LXR agonist, GW3965, incorporated into synthetic high-density lipoprotein nanodiscs (sHDLs). sHDLs are 8-12 nm in size and are composed of apolipoprotein mimetic peptides, in this case, 22A,^[19]^ complexed with phospholipids. sHDLs mimic naturally occurring pre-β HDL which contains apolipoprotein A-I, phospholipids, and unesterified cholesterol in the body, and can remove cholesterol and phospholipids from cells through their interactions with the ABCA1 transporter.^[20]^ Because of this, sHDLs are an attractive nanocarrier system for LXR agonists due to the inherent synergy wherein sHDL can act as an acceptor of excess cholesterol that accumulates within GBM tumor cells. In addition, a growing body of research from our lab and other investigators demonstrates the therapeutic efficacy of sHDLs delivering LXR agonists in other diseases such as atherosclerosis.^[21–23]^ Our lab has also previously shown that sHDL-mediated delivery of CpG and docetaxel improved survival in GBM tumor-bearing mice.^[19, 24, 25]^ Furthermore, sHDL nanodiscs have been administered to humans in Phase I/II clinical trials, so there is established evidence of safety, which is advantageous for translating sHDL-based therapies into the clinic.^[19, 26, 27]^

Our working hypothesis in this study is that treatment with LXR agonists and CpG delivered by sHDL combined with standard radiation therapy would function as a more effective treatment strategy in GBM tumor-bearing mice. To test this hypothesis, we treated glioma-bearing mice over the course of several weeks with drug-loaded sHDL, free LXR agonist, radiation therapy alone, or drug-loaded sHDL in combination with radiation therapy. Following the treatment period, mice were rechallenged with tumor cells after 90 days to evaluate changes in immune protection against tumor growth.

## 2. Results

### Preparation and characterization of sHDL Nanodiscs containing LXR Agonists and CpG

Liver X Receptor (LXR) agonists have been investigated extensively in targeting cholesterol accumulation, primarily in the context of atherosclerosis. Given the dependency of glioblastoma (GBM) tumors on cholesterol produced by neighboring astrocytes and the immunosuppression caused by cholesterol accumulation in the tumor microenvironment, we prepared LXR agonist-loaded synthetic high-density lipoprotein (sHDL) nanodiscs. sHDL represents an attractive carrier due to its inherent synergy with LXR agonists, wherein sHDL can act as an acceptor of cholesterol effluxed by target cells through direct interaction with the ABCA1 transporter. In addition, the ability of sHDL to deliver CpG antigens to immune cells and more efficiently traffic through tumors due to its small size (8-12 nm) has been previously demonstrated,^[24, 25, 28]^ further supporting the use of sHDL as a drug carrier.

In order to determine the effectiveness of sHDL-delivering LXR agonists, we first synthesized nanodiscs with cholesterol-modified CpG 1826 (chol-CpG) and 1 of 3 different LXR agonists (GW3965 [GW], RGX104 [RGX], or T0901317 [T0]) incorporated into the membrane. sHDL nanodiscs were prepared as previously described using a co-lyophilization method in which the apolipoprotein mimetic peptide 22A, DMPC, and LXR agonist were dissolved in acetic acid and freeze-dried overnight (Figure 1A). The resulting lyophilized powder was then rehydrated in pH 7.4 PBS, followed by heating at 50°C and cooling on ice 3 times (above and below the transition temperature of DMPC) to facilitate final sHDL assembly. The encapsulation efficiency for all 3 drugs was >90% (final drug concentration ∼ 1.5mg/mL) as determined by UPLC-MS following the separation of Shdl encapsulated LXR agonists from free drug using a 7000k MWCO desalting column. Chol-CpG was inserted into purified nanodiscs through the cholesterol moiety by incubation with LXR agonist encapsulated-sHDL in PBS for 1 hour at room temperature to yield the final LXR agonist and CpG co-encapsulated sHDL nanodiscs (LXR-CpG-sHDL).

**Figure 1.**
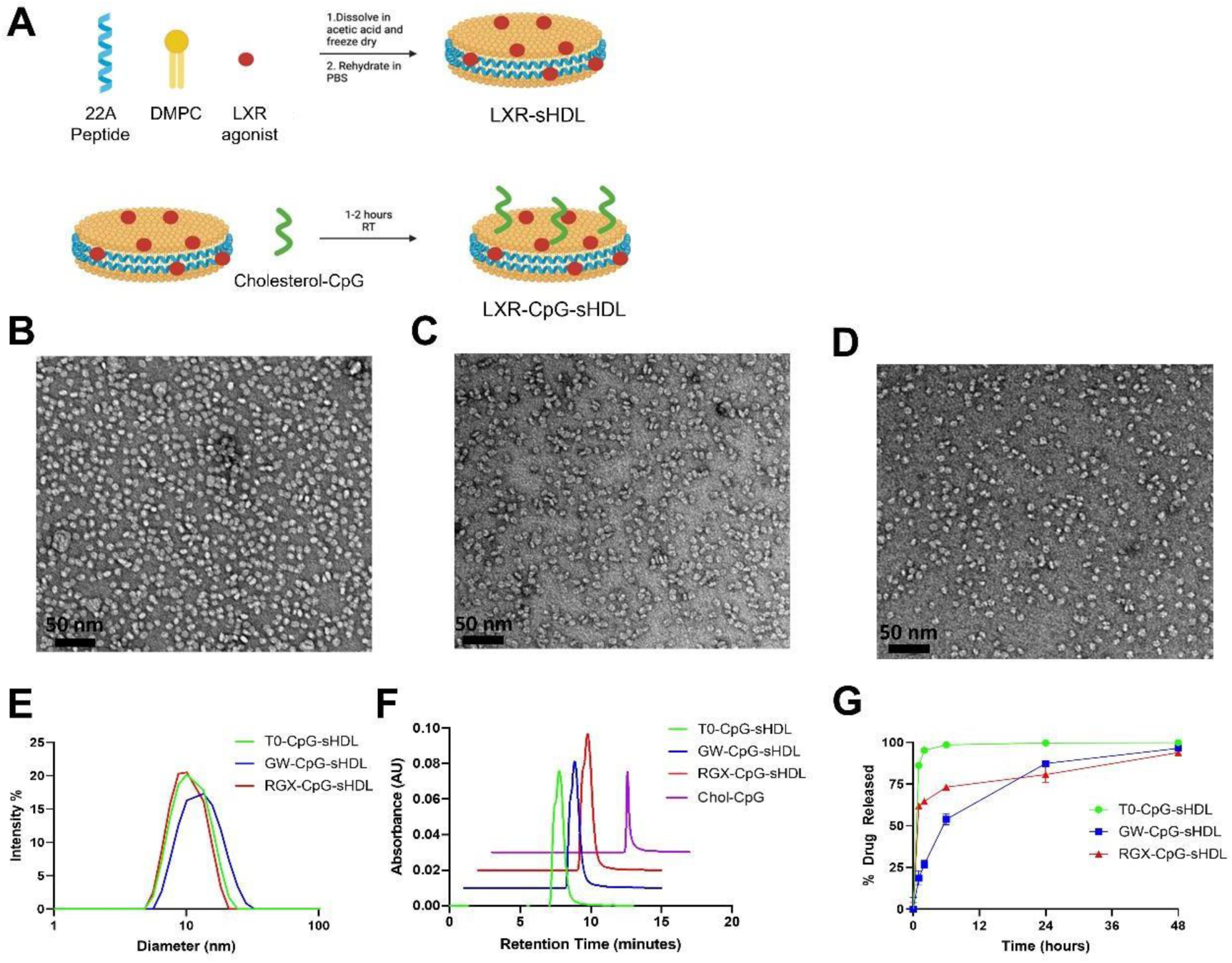
Characterization of LXR agonist encapsulated sHDL-CpG nanodiscs (LXR-CpG-sHDL). Preparation workflow for LXR-sHDL-CpG nanodiscs (A). Negative-stain TEM images of T0 (B), GW (C), and RGX (D) encapsulated nanodiscs. Particle size distribution of LXR-CpG-sHDL formulations determined via DLS (E). Purity of nanodiscs determined using size exclusion chromatography with absorbance at 260 nm (F). Release profiles of LXR agonists from sHDL particles in PBS containing BioBeads adsorbent media (G). N=3, Mean ± SD.

Following sHDL synthesis, the morphology of each formulation was observed using negative-stain TEM (Figure 1B-D). The TEM images revealed a similar disc-like morphology and particle size for each formulation. Nanodisc size and homogeneity were further analyzed using dynamic light scattering (DLS). All 3 formulations were 10-12nm in diameter with a polydispersity index of ∼0.2 (Figure 1E). Analysis of each formulation via size-exclusion chromatography revealed complete insertion of chol-CpG into the sHDL membrane, as no free chol-CpG was detected following the incubation (Figure 1F). The successful insertion of chol-CpG was also evidenced by the reduction of surface zeta potential after chol-CpG insertion due to the anionic nature of CpG moiety (Figure S1). In order to compare the stability of each LXR agonist within the sHDL membrane, the release profile of all 3 LXR agonists from sHDL nanodiscs was determined in PBS containing BioBeads adsorbent media at 37°C. All 3 drugs showed complete release from the nanodisc within 48 hours, with T0 having the most rapid release profile, followed by more gradual release profiles for RGX and GW (Figure 1G). Visual inspection of the T0-CpG-sHDL formulation during the incubation revealed precipitation of free T0, which explains the nearly complete release observed after 30 minutes.

### *In Vitro* Efficacy of LXR-CpG-sHDL

The mechanism of action of LXR agonists has been well-described in the literature, with their applications ranging from atherosclerosis to cancer.^[29]^ LXRs are nuclear receptors that, when activated by LXR agonists, for heterodimers with Retinoid-X-Receptors (RXRs) to regulate the expression of genes with downstream effects ranging from cholesterol efflux to suppression of inflammation and immune cell modulation. After synthesizing the 3 different LXR-CpG-sHDL, we next compared their impact on cholesterol-transport gene expression, cholesterol efflux, and cytotoxicity in mouse glioma cells. Among the downstream genes affected by LXR activation, we chose to analyze LXRα, LXRβ, and ABCA1, with our primary interest being ABCA1 due to its role in cholesterol efflux and loading of endogenous HDL in the body. qPCR analysis following overnight treatment with 5 µM LXR agonist (either free or incorporated into sHDL) did not have a substantial impact on the mRNA levels of LXRα or LXRβ (Figure 2A-B), as only free RGX and RGX nanodiscs resulted in a minor (∼1.2-fold) increase in LXRβ mRNA.

**Figure 2.**
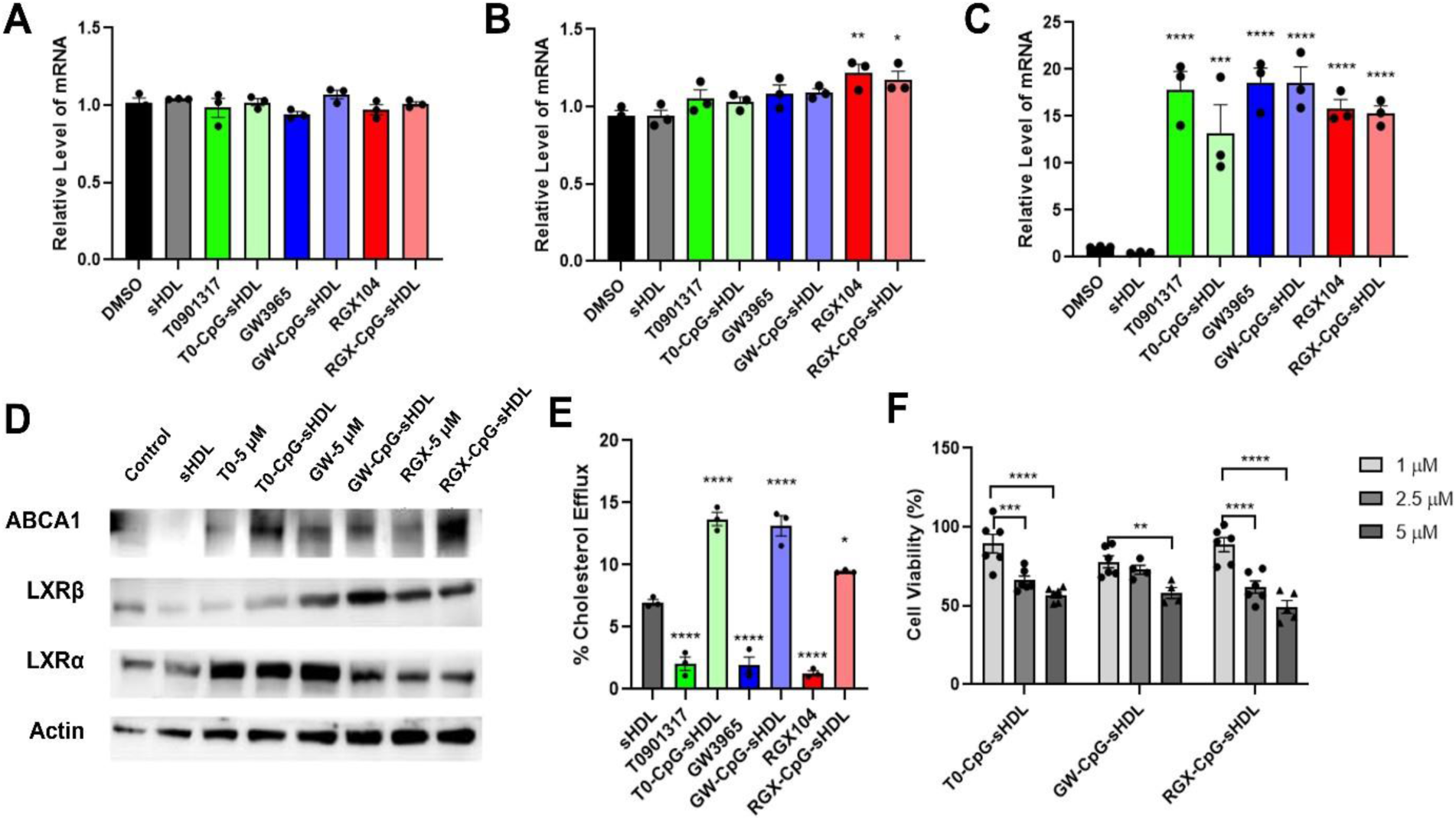
Efficacy of free LXR agonists and LXR-CpG-sHDL nanodiscs in GL26 cells. Changes in mRNA levels of LXRα (A), LXRβ (B), and ABCA1 (C) following treatment with 5 µM free or sHDL-loaded LXR agonists evaluated via qPCR. Changes in protein expression evaluated via western blot (D). Radiolabeled cholesterol efflux from GL261 cells following 24-hour treatment with 1 µM LXR agonist or sHDL formulation (E). Viability of GL261 cells at increasing concentrations of LXR-CpG-sHDL formulations. *p<0.05, **p<0.01, ***p<0.001, ****p<0.0001. N=3, Mean ± SEM.

In contrast, all LXR agonist treatments resulted in significant increases in the mRNA levels of ABCA1 (Figure 2C), with free GW and GW encapsulated nanodiscs resulting in the most significant change (∼20-fold increase). Subsequent analysis at the protein level by western blot revealed modest increases in LXRα levels for each of the LXR treatments relative to DMSO (control) or blank sHDL-treated cells (Figure 2D). In addition, only GW and RGX (free drug and sHDL formulations) increased the amount of LXRβ present. Consistent with the qPCR data, all LXR treatments resulted in an increase in ABCA1.

After confirming the LXR-loaded nanodiscs are able to increase expression of the ABCA1 cholesterol efflux transporter, we compared the effectiveness of each treatment with regards to cholesterol removal from glioma cells, as this is expected to be one of the main drivers of tumor cell death *in vivo*. To measure cholesterol efflux, cells were loaded with radiolabeled cholesterol [1,2-^3^H(N)] followed by 24-hour treatment with DMSO, blank sHDL (no LXR agonist), free LXR agonist, or LXR agonist encapsulated sHDL nanodiscs (Figure 2E). As expected, given its known role as a cholesterol acceptor, treatment with blank sHDL resulted in modest (∼7%) cholesterol removal from the cells. Treatment with free LXR agonists had a minimal effect (1-2%) on cholesterol removal, which is to be expected given the lack of any lipoprotein cholesterol acceptor present. Importantly, all 3 sHDL formulations of the LXR agonists showed significant improvements in cholesterol removal compared to blank sHDL, with T0- and GW-encapsulated sHDL nanodiscs showing the highest % cholesterol efflux (∼14%).

Each sHDL formulation was next compared in its ability to induce cell death in glioma cells at increasing LXR agonist doses (Figure 2F). All 3 LXR-CpG-sHDL formulations reduced cell viability to a similar degree in a dose-dependent manner at concentrations up to 5 µM (∼40% reduction in cell viability). Based on the superior membrane retention/stability of the GW-CpG-sHDL formulation and its high cholesterol efflux, the GW-CpG-sHDL nanodiscs were selected for further investigation in human and mouse high-grade glioma cells and in glioma tumor-bearing mice. To assess the efficacy of GW-CpG-sHDL nanodiscs in inducing cell death and enhancing radio-sensitivity in high-grade gliomas, we conducted cell viability assays using mouse GL-26 EGFRvIII cells and two human patient-derived GBM cells, HF2303 and MGG8, in conjunction with irradiation. We used normal human astrocytes as control cells to assess any potential undesired cytotoxic effects of the nanodisc treatment. Cells were exposed to saline, free-sHDL, CpG-sHDL, free-GW3965, and GW-CpG-sHDL nanodiscs with or without irradiation. Our findings indicate that treatment with increasing doses of either free-GW3965 or GW-CpG-sHDL nanodiscs alone significantly inhibited cell survival in a dose-dependent manner at concentrations up to 5 µM in both mouse (Figure 3B-C) and human glioma cells (Figure 3D-G). Moreover, treatment with GW-CpG-sHDL nanodiscs consistently reduced the survival of glioma cells tested, with reductions ranging from 10% to 65%, 28% to 68%, and 16% to 31%, compared to saline-treated controls for GL-26 EGFRvIII, HF2303 and MGG8 cells, respectively. In contrast, normal human astrocytes showed minimal and biologically insignificant cytotoxic effects following treatment with GW3965, GW-CpG-sHDL, or empty-nanodisc (Figure S2). Notably, combined treatment with free-GW3965 or GW-CpG-sHDL nanodiscs and irradiation increased radio-sensitivity, resulting in a greater decrease in cell viability across all GBM cell lines tested compared to treatment with free-sHDL, CpG-sHDL, irradiation alone, or their combinations (Figure 3).

**Figure 3.**
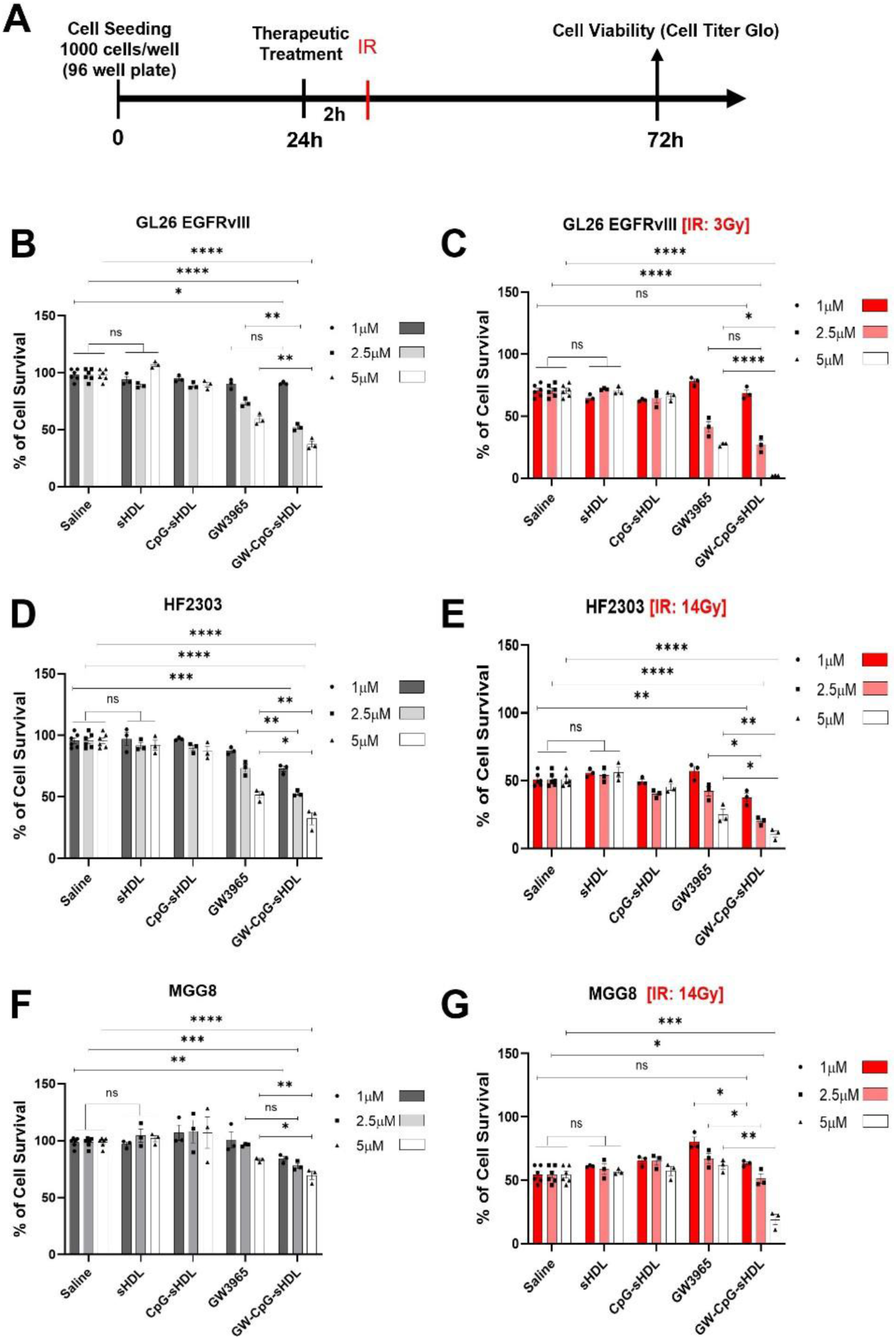
*In vitro* activation of Liver X Receptor (LXR) and TLR9 through free GW3965 (LXR agonists), CpG-sHDL (TLR9 ligand loaded nanodiscs) and GW-CpG-sHDL nanodiscs promotes cytotoxicity and enhances radiosensitivity in murine and human patient-derived glioblastoma cells. (A) Schematic illustration shows the timeline of the *in vitro* application of free-GW3965 and GW-CpG-sHDL nanodiscs in combination with radiation in mouse-glioblastoma (GL26 EGFRvIII) and patient-derived glioblastoma cells (MGG8 and HF2303). (B-C) GL26 EGFRvIII, (D-E) HF2303 and (F-G) MGG8 cells were treated with either free-GW3965, empty-sHDL, CpG-sHDL and GW-CpG-sHDL nanodiscs alone or in combination with radiation (3Gy for mouse cells and 14Gy for human cells) at 1, 2.5 or 5 µM doses for 72 h. sHDLs without GW3965 were tested in the same peptide and lipid concentrations as GW-CpG-sHDL with specified GW3965 concentration. Bars represent mean ± SEM (n = 3 biological replicates). ns= not significant, *p<0.05 **p< 0.01, ***p< 0.0001, ****p< 0.0001.

### GW-CpG-sHDL induces caspase3/7 dependent apoptotic cell death in high-grade gliomas

Reports disclosed that different LXR agonists can promote cell death either by inhibiting LDLR expression or by enhancing the anti-proliferative effect of BH3 mimetics in different solid tumor models^[12, 30–32]^. Additionally, these agonists can reduce the suppressive functions of MDSCs on T cells by promoting apoptosis of MDSCs within the tumor microenvironment^[32]^. To explore whether GW-CpG-sHDL induced cell death involves any cell cycle phases, potentially corroborating apoptotic cell death in glioblastoma, we performed cell cycle analysis using propidium iodide (Figure S3). Flow cytometry revealed a significant increase in the sub-diploidal (sub-G0/G1) population in both GL26 EGFRvIII and HF2303 glioblastoma cells after 72 hours of treatment with GW3965 and GW-CpG-sHDL nanodiscs (Figure 4A-B). Specifically, the sub-diploidal population increased to 18.37% and 33.6% for GL26 EGFRvIII and 23.77% and 34.93% for HF2303 with GW3965 and GW-CpG-sHDL, respectively, compared to untreated or empty-nanodisc controls (GL26 EGFRvIII: 1.21%, 1.23%; HF2303: 0.85%, 1.29%, respectively). The increase was significantly greater with GW-CpG-sHDL compared to GW3965 alone. Further analysis using the human patient-derived glioblastoma cells MGG8 under similar conditions confirmed this trend, with the sub-diploidal population increasing to 13.1% and 22.73% for GW3965 and GW-CpG-sHDL, respectively, compared to 1.90% and 3.21% for untreated and empty-nanodisc controls (Figure S4A). Again, GW-CpG-sHDL showed a significantly higher effect that GW3965 alone. In contrast, normal human astrocytes exhibited only minimal and biologically insignificant changes in sub-diploidal populations (3.15%, 4.47%, 5.01% and 3.57% for untreated, GW3965, GW-CpG-sHDL, and empty-nanodisc treatments, respectively (Figure S4B). Staurosporine-treated cells served as positive controls. These findings indicate that GW-CpG-sHDL selectively increases sub-diploidal populations in glioblastoma cells, suggesting apoptotic activity, while sparing normal human astrocytes. This underscores its potential for targeted therapeutic strategies in glioblastoma.

**Figure 4.**
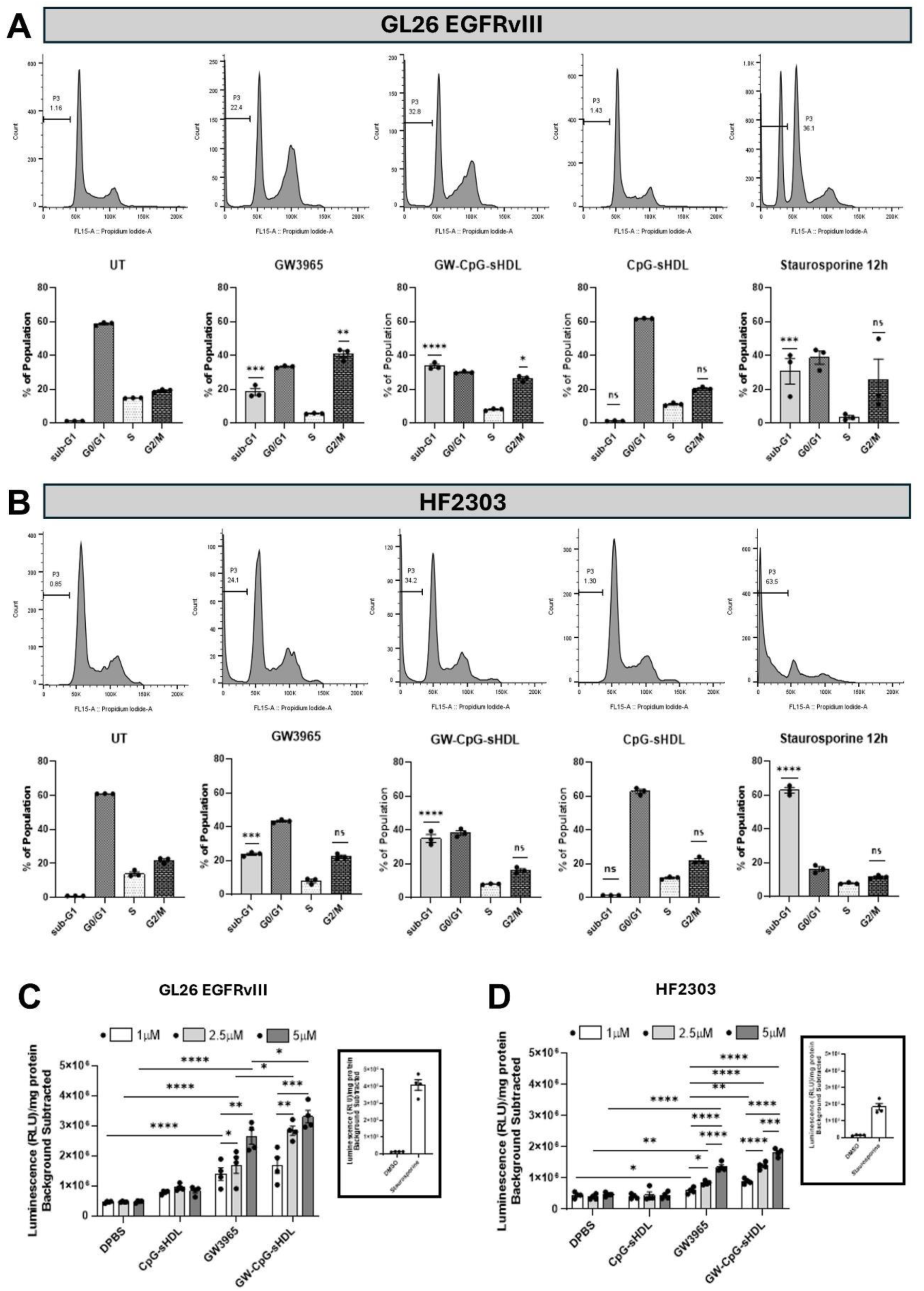
Cell cycle distribution and caspase 3/7 activation in mouse and human glioblastoma cells following treatment with LXR agonist and LXR agonist-loaded nanoparticles. (A) Mouse GL26 EGFRvIII and (B) human HF2303 cells were treated with either free-GW3965, empty sHDL, CpG-HDL, or GW-CpG-sHDL nanodiscs at a concentration of 5 µM for 72 hours. Staurosporine (0.5 µM) served as a positive control for 12 hours. sHDLs without GW3965 were tested using the same peptide and lipid concentrations as those used in GW-CpG-sHDL treatments, with corresponding GW3965 doses. Cells were harvested and fixed in 70% ethanol, stained with propidium iodide, and analyzed via flow cytometry. The percentage of cells in the sub-G1 phase (indicative of hypodiploid DNA content), is shown in each panel. Bar graphs display the “% of population” in different cell cycle phases, with statistical significance of other treatment groups compared to the untreated control. To assess caspase 3/7 activation, (C) GL26 EGFRvIII and (D) HF2303 cells were treated with either free-GW3965, empty-sHDL, CpG-HDL, or GW-CpG-sHDL nanodiscs at concentrations of 1, 2.5, or 5 µM for 12 hours. The Caspase–Glo 3/7 apoptosis assay showed a significant increase in normalized luminescence values in treated groups compared to the untreated control, indicating pro-apoptotic bioactivity in a dose-dependent manner. Staurosporine (0.5 µM) and DMSO (20%) were used as positive and negative controls, respectively, for 12 hours (data presented in the inset). Values represent the mean ± SEM of three independent experiments with four replicates per condition. Data were analyzed using one-way ANOVA followed by post hoc Tukey’s test and unpaired Student’s t-test to assess significant differences between groups. Statistical significance relative to the untreated control is indicated as *p < 0.05, **p < 0.01, ***p < 0.001, ****p < 0.0001.

Apoptosis can be triggered by several factors, leading to the activation of executioner caspases. Among these, caspase 3/7 death proteases are indispensable and are frequently activated during drug treatment, driving cells towards programmed cell death ^[33, 34]^. To investigate whether caspase 3/7 activation contributes to GW-CpG-sHDL induced cell death in mouse and human glioblastoma cells, we performed a luminescence-based caspase-3/7 activation assay on mouse glioblastoma, human glioblastoma, and normal human astrocytes (Figure S4C). We found that GW3965 and GW-CpG-sHDL significantly increased the caspase-3/7 activity in GL26 EGFRvIII (Figure 4C) and HF2303 (Figure 4D) glioblastoma cells at 12 hours, compared to empty-nanodisc treated or untreated cells. Moreover, GW-CpG-sHDL elicited significantly higher caspase-3/7 activity than GW3965 alone in both GL26 EGFRvIII and HF2303 cells (Figure 4C-D). Similarly, in human glioblastoma MGG8 cells treated under identical conditions, caspase-3/7 activity was significantly elevated after treatment with GW3965 and GW-CpG-sHDL, with the highest activity observed in GW-CpG-sHDL treated cells (Figure 4D). In contrast, caspase-3/7 activity remained low and biologically insignificant in normal human astrocytes treated with the same nanodiscs (Figure S4E). Staurosporine and DMSO-treated cells served as positive and negative controls, respectively, in all experimental groups. These findings confirms that GW-CpG-sHDL-induced cell death involves caspase-3/7 in both mouse and human glioblastoma cells, while sparing the normal human astrocytes.

### Post-Operative Recurrence of GBM Necessitates Intracranial Administration

Developing effective treatments for patients with Glioblastoma Multiforme (GBM) continues to be a challenge in clinical neuro-oncology. The current standard of care for GBM patients involves surgical resection of the tumor mass, complemented by radiation therapy and adjuvant temozolomide (TMZ). However, due to the invasive nature of gliomas and their proximity to critical brain structures, complete tumor removal is not always feasible. Factors such as aggressive tumor behavior or the presence of residual disease increase the risk of early recurrence. The site of glioma recurrence is typically concentrated around the boundaries of the resected area (Figure 5). Therefore, it is crucial to inject therapeutic agents directly along these margins to minimize the risk of tumor recurrence. By targeting the peripheral regions of the resected tumor site, we can ensure that residual tumor cells, which are often responsible for recurrence, are effectively treated. This localized delivery approach enhances the concentration of the therapeutic agent where it is most needed, thereby improving the chances of long-term remission and reducing the likelihood of the tumor returning.

**Figure 5.**
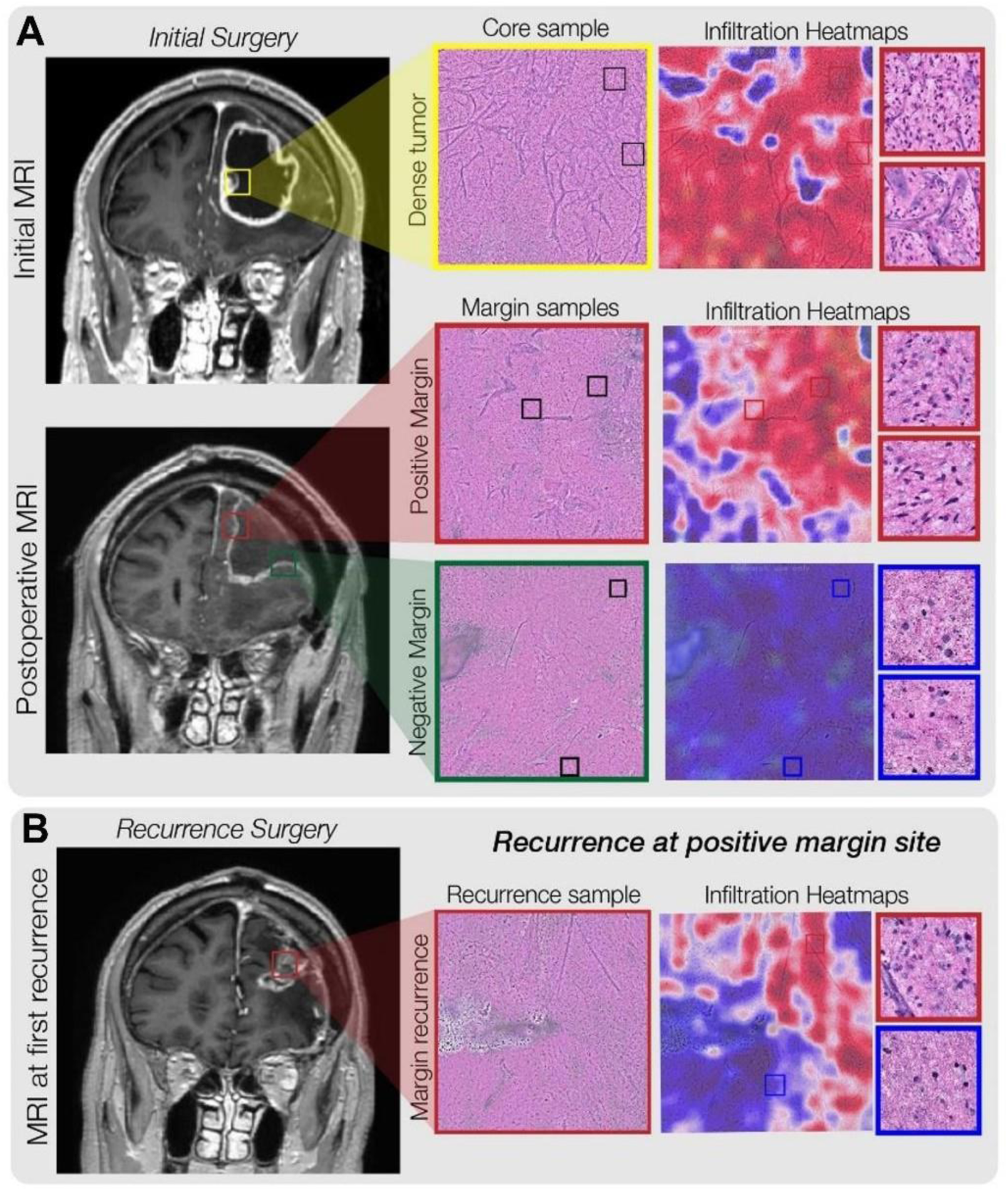
An illustrative example of predicting tumor recurrence using intraoperative optical imaging and machine learning models to detected and quantify diffuse glioma infiltration. a, Intraoperative stimulated Raman histology (SRH) is used to image the surgical margin of diffuse glioma resection cavities. Within fresh, unprocessed, unstained surgical specimens, tumor infiltration can be identified using transformer-based deep neural networks. Our SRH models provide interpretable visualizations to assess the degree of tumor infiltration within surgical specimens from diffuse glioma patients. Top, a tumor specimen is sampled from the contrast enhancing core of a glioblastoma. Dense tumor is identified using our SRH tumor infiltration heatmaps. Middle, specimens were sampled from the medial and lateral margins of the resection cavity at completion of the operation. No tumor infiltration was identified at the lateral surgical margin; however, sparse tumor infiltration was noted within the white matter at the medial surgical margin at completion of the surgery. b, MRI of the same patient at first evidence of radiographic recurrence. A nodule of contrast enhancing tissue was noted on the medial aspect of the resection cavity. Intraoperatively, dense recurrence identified on SRH was noted within the surgical specimen sampled from this area of radiographic recurrence. Our SRH tumor infiltration model identified definitive tumor infiltration within the specimen.

### Efficacy of GW-CpG-sHDL nanodiscs in Glioma Bearing Mice

To fully elucidate the combination effects of GW3965 and CpG, we conducted experiments with the following treatment groups: saline (median survival [MS] = 25 days), sHDL (MS = 26 days), GW3965+CpG (MS = 27 days), sHDL-CpG+IR (MS = 30 days), and GW3965+CpG+IR (MS = 38 days) (Figure S5). The sHDL group showed a marginal increase in median survival compared to saline, while the GW3965+CpG group demonstrated a slight improvement. Notably, the combination of sHDL-CpG with IR significantly enhanced median survival compared to saline. The most substantial improvement was observed in the GW3965+CpG+IR group, which showed a median survival of 38 days. This was the only group that showed a significant improvement in median survival compared to saline, indicating a significant synergistic therapeutic effect. These findings underscore the enhanced efficacy of combining GW3965, CpG, and IR, providing substantial survival benefits and supporting their potential as a robust therapeutic strategy for glioblastoma. To assess the efficacy of GW-CpG-sHDL nanodiscs *in vivo*, GBM-bearing mice were treated with 4 doses of saline, free GW3965, or GW-CpG-sHDL nanodiscs over a 2-week treatment period (Figure 6A). Since radiation therapy is the current standard of care for GBM, we tested the efficacy of free GW3965 and GW-CpG-sHDL alone or in combination with two cycles of radiotherapy on the survival of tumor-bearing mice. The results showed that treatment with IR alone increased the median survival (MS) to 34 days compared to the saline group MS=28 (Figure 6B). Administration of free GW3965 or GW-CpG-sHDL increased the median survival to 34 days and 33 days, respectively, with GW-CpG-sHDL treatment resulting in 1 long-term survivor. Combining GW3965 treatment with IR showed negligible improvement compared to either treatment alone (MS=35). Interestingly, treatment with GW-CpG-sHDL in combination with IR resulted in substantial improvements in survival compared to either treatment alone (MS=not reached), with 60% of the mice surviving long-term.

**Figure 6.**
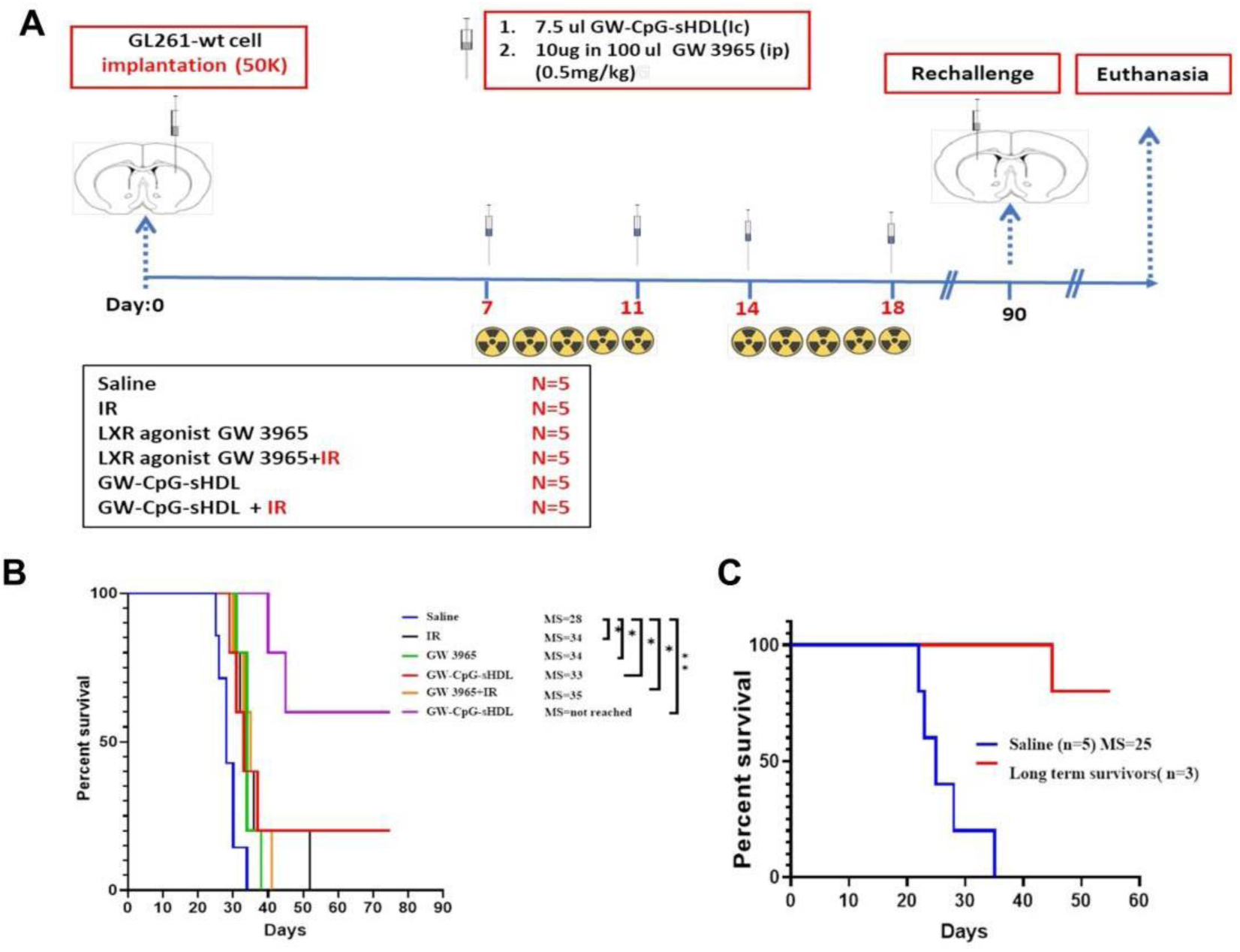
Efficacy of GW-CpG-sHDL nanodiscs is GL261-tumor bearing mice. Outline of treatment schedule (A) with mice receiving saline, IR, free GW3965, GW-CpG-sHDL, GW3965 + IR, or GW-CpG-sHDL + IR at the indicated timepoints. Kaplan-Meier survival curves for each treatment group (B). Survival curve of rechallenged survivors from the GW-CpG-sHDL + IR group (C). N=5 mice per group. *p<0.05, **p<0.01.

The long-term survivors from the GW-sHDL-CpG + IR treatment group were rechallenged with glioma cells implanted in the contralateral hemisphere. They received no additional treatment in order to assess the development of immunological memory. Compared to saline-treated mice (MS=25), the rechallenged survivors showed significantly improved survival, with 2 of the 3 mice showing no tumor after the rechallenge (Figure 6C).

Subsequent analysis of brain sections from long-term survivors showed no evidence of intracranial tumor after 60 days of tumor cell implantations. (Figure 7). In contrast, hematoxylin and eosin (H&E) staining of saline-treated mice showed the presence of the tumor. In the long-term survivors who underwent the combination therapy, no discernible regions exhibiting hemorrhages, necrosis, or invasion were observed (Figure 7). To evaluate the potential impact of the combination treatment on the surrounding brain architecture, immunohistochemical (IHC) staining was performed employing glial fibrillary acidic protein (GFAP) and myelin basic protein (MBP) as markers for assessing the integrity of the myelin sheath. The results demonstrated no apparent alterations in brain architecture among the mice subjected to the combined GW-CpG-sHDL+IR treatment compared to the control group (Figure 7).

**Figure 7.**
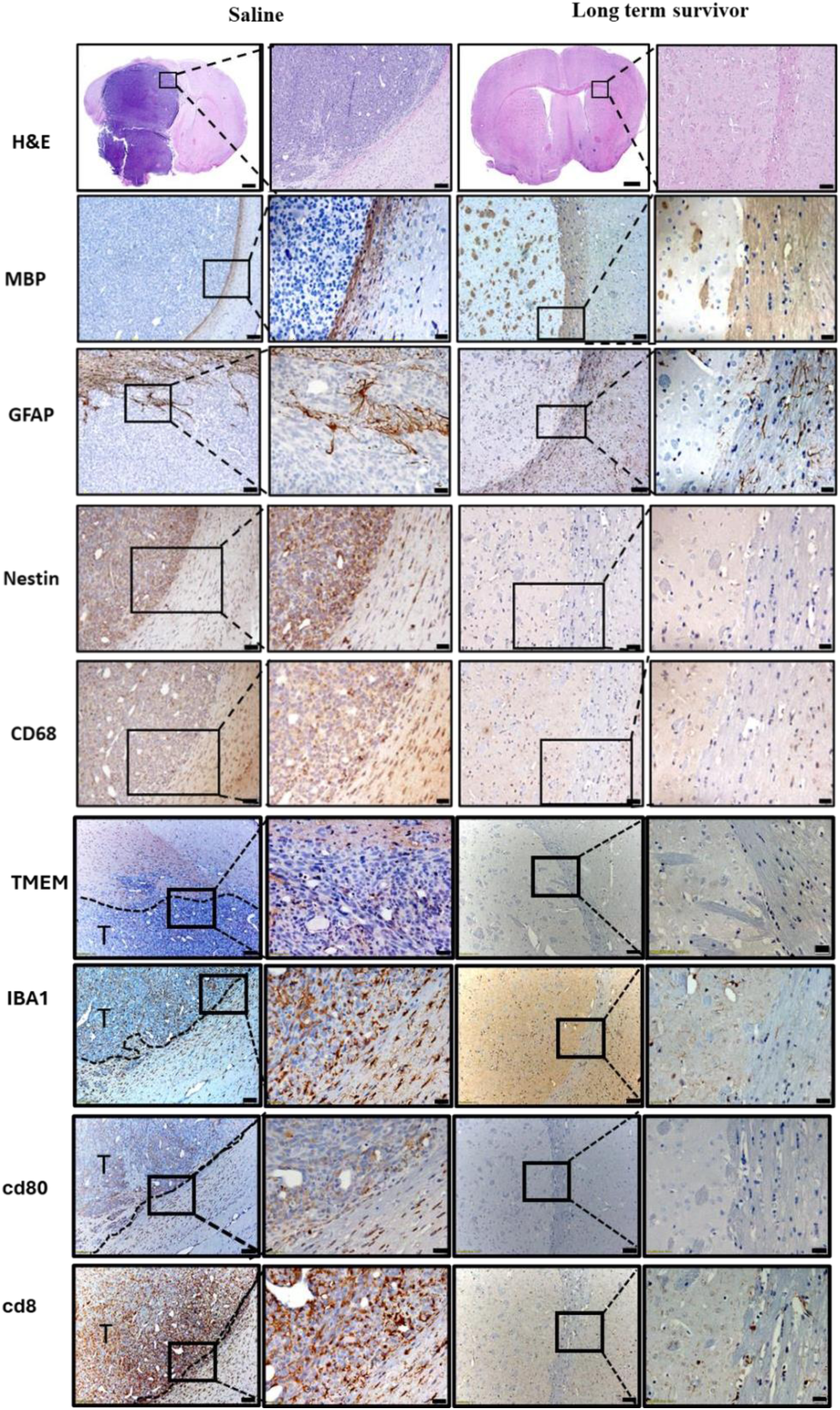
H&E and Immunohistochemistry staining of 5µm paraffin-embedded brain sections from saline treated mice and long-term survivors from GW-CpG-sHDL treated mice. The combination therapy group exhibits reduced immune cell activation and infiltration, indicating a modulated immune response and contributing to therapeutic success and long-term survival. Low magnification (black scale bar = 50um for nestin and CD68, 100 μm for other staining images). High magnification (black scale bar = 20µm). Representative images from a single experiment consisting of independent biological replicates are displayed.

Immunohistochemical analysis was conducted to evaluate the immune landscape within the brain following treatment, employing markers TMEM119, IBA1, CD80, and CD8. TMEM119, indicative of microglia, showed increased presence in the tumor microenvironment and adjacent brain tissue of saline-treated mice, signifying heightened microglial activation (Figure 7). Conversely, the combination therapy group exhibited fewer TMEM119+ cells, suggesting reduced microglial activation and a less inflammatory environment. IBA1, a marker for microglia and macrophages, was more prevalent in saline-treated mice, indicating elevated microglial and macrophage activity, whereas the combination therapy group showed reduced IBA1+ cells, correlating with diminished inflammation and tumor burden. CD80, a co-stimulatory molecule on activated antigen-presenting cells, was more abundant in the tumor microenvironment of saline-treated mice, reflecting active immune response, while the long-term survivors from combination therapy group had fewer CD80+ cells, indicating decreased immune activation due to effective tumor control. CD8, a marker for cytotoxic T lymphocytes, was present in saline-treated mice, indicating an immune attempt to combat the tumor, whereas long term long-term survivors from combination therapy exhibited lower CD8+ T cells. Collectively, these findings indicate that the combination therapy modulates immune cell presence and activity, reducing microglial and macrophage infiltration (TMEM119 and IBA1), maintaining effective immune surveillance (CD8), and reducing unnecessary immune activation (CD80). This balance likely contributes to the observed therapeutic success and long-term survival in treated mice.

Additionally, we conducted the immunohistochemical (IHC) analysis of brain tumor sections from control and treated mice at day 30 post-tumor implantation;12Days after the last treatment time point study. Figure 8 reveals significant differences in tumor morphology (Figure 8B and C), immune response (Figure 8D and E), and cellular markers (Figure S6B and C) between control and treated brain tumor sections. H&E staining of brain tumor sections shows larger, more invasive tumor regions in the control group compared to smaller, well-defined tumor boundaries in treated mice (Figure 8B and C). CD8+ T-cell staining reveals markedly increased cytotoxic T-cell infiltration in treated tumor sections compared to controls, highlighting the activation of an anti-tumor cytotoxic immune response. Similarly, CD4+ T-cell staining shows elevated helper T-cell presence in treated tumors, supporting the role of adaptive immunity in enhancing treatment efficacy (Figure 8B and C). The percentage of CD4+ and CD8+ cell percentages in brain tumor sections from control and treated mice at day 30 post-tumor implantation; 12 days after the last treatment was also assessed, Both CD4+ (p-value: 0.00026) and CD8+ (p-value: 0.000078) cell percentages were found significantly higher in the treated group compared to the saline group (Figure 8D and E). Myelin basic protein (MBP) staining shows extensive myelin loss in control tumors, while treated sections exhibit better preservation of myelin integrity, indicating reduced tumor-associated damage. Glial fibrillary acidic protein (GFAP) staining indicates higher astrocyte activation at tumor margins in control sections, while treated tumors show reduced glial reactivity. Iba1 staining highlights dense microglial activation and infiltration in control tumors, which is significantly diminished in treated sections (Figure S6B and C).

**Figure 8.**
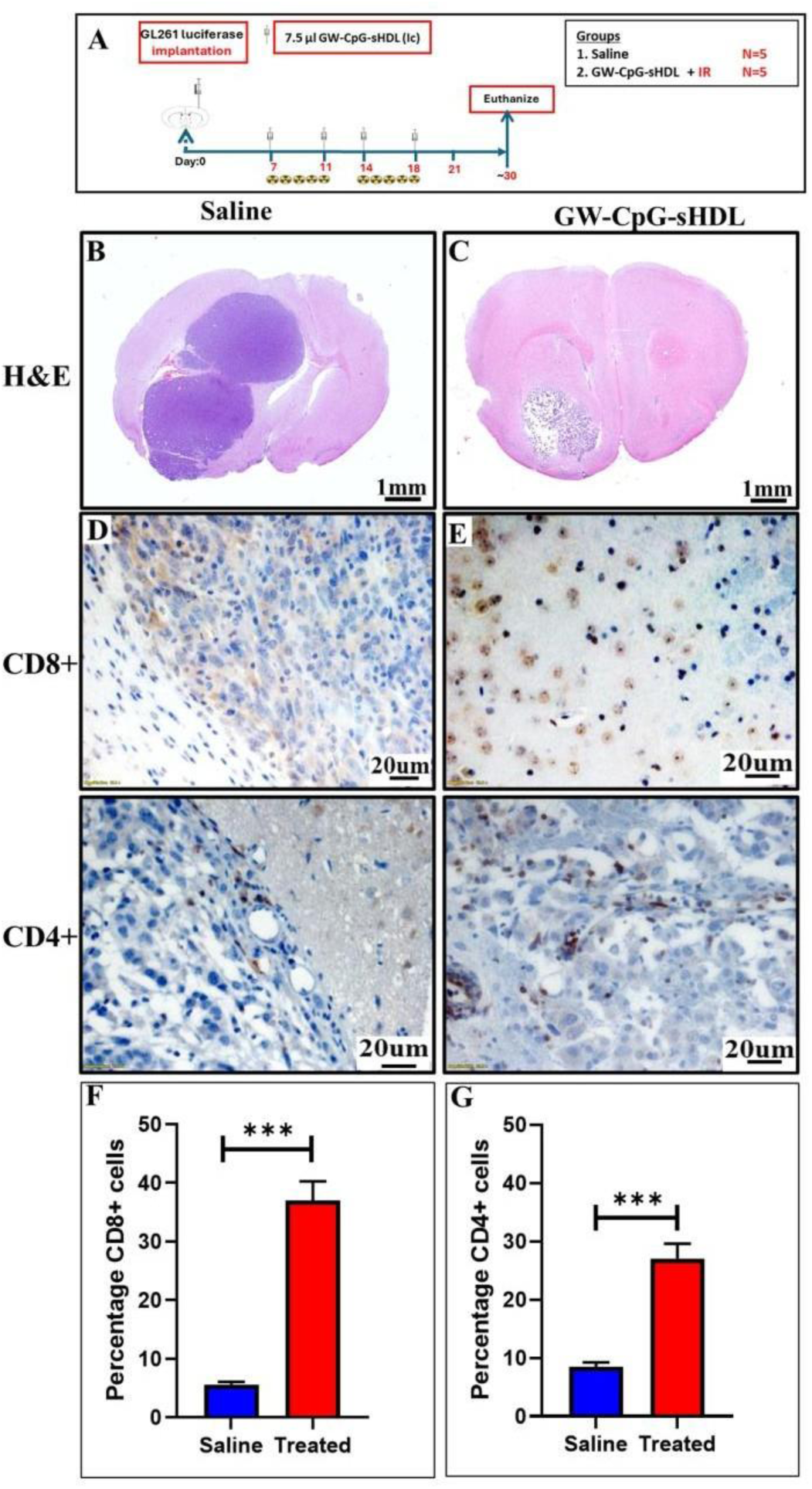
Immunohistochemical analysis of brain tumor sections from control and treated mice at day 30 post-tumor implantation;12Days after the last treatment. (A) Outline of treatment schedule for the 30-day timed experiment. (B, C) H&E staining shows reduced tumor burden in treated mice compared to extensive tumor growth in controls. CD8+ T-cell staining reveals markedly increased cytotoxic T-cell infiltration in treated tumor sections (D, E). Similarly, CD4+ T-cell staining shows elevated helper T-cell presence in treated tumors. Bar graph showing the percentages of CD8+ (left) (F) and CD4+ (right) (G) cells in the saline (blue) and treated (red) groups. Both CD4+ (p-value: 0.00026) and CD8+ (p-value: 0.000078) cell percentages are significantly higher in the treated group compared to the saline group.

We also evaluated a panel of biomarkers for each treatment group’s healthy liver and kidney function from the initial survival study (Figure S7). Serum levels of creatine, BUN, ALT, ALB, ALKP, and TPRO did not significantly change as a result of any of the treatments. In addition, H&E staining of liver sections from saline-treated mice and long-term survivors revealed no adverse treatment effects on the liver (Figure S8).

These data demonstrate that treatment with GW-CpG-sHDL combined with IR can elicit a potent antitumor response, leading to long-term survival and immunological memory with no measurable adverse effects on the brain tissue or major peripheral organs.

### Tumor progression and bioluminescence imaging

The results show the therapeutic efficacy of GW-sHDL-CpG combined with irradiation (IR) in reducing tumor progression in tumor-bearing mice compared to saline-treated controls. Bioluminescence imaging (BLI) showed a consistent increase in tumor radiance over time in the saline group, indicating unchecked tumor growth. In contrast, mice treated with GW-sHDL-CpG + IR exhibited significantly reduced tumor radiance, suggesting effective tumor suppression. This reduction in tumor burden was observed consistently across all treated mice, as evident from the lower intensity of bioluminescent signals compared to saline-treated animals (Figure 9B and C). Radiance plots revealed a sharp increase in radiance over time in the saline group, reflecting aggressive tumor growth(Figure 9D and E). Conversely, mice in the GW-sHDL-CpG + IR group exhibited a peak in tumor radiance, followed by a significant decline, indicating a reduction in tumor viability and effective tumor control (Figure 9). These findings highlight the potential of the GW-CpG-sHDL + IR combination therapy to suppress tumor progression, with a marked difference in tumor dynamics compared to untreated controls.

**Figure 9.**
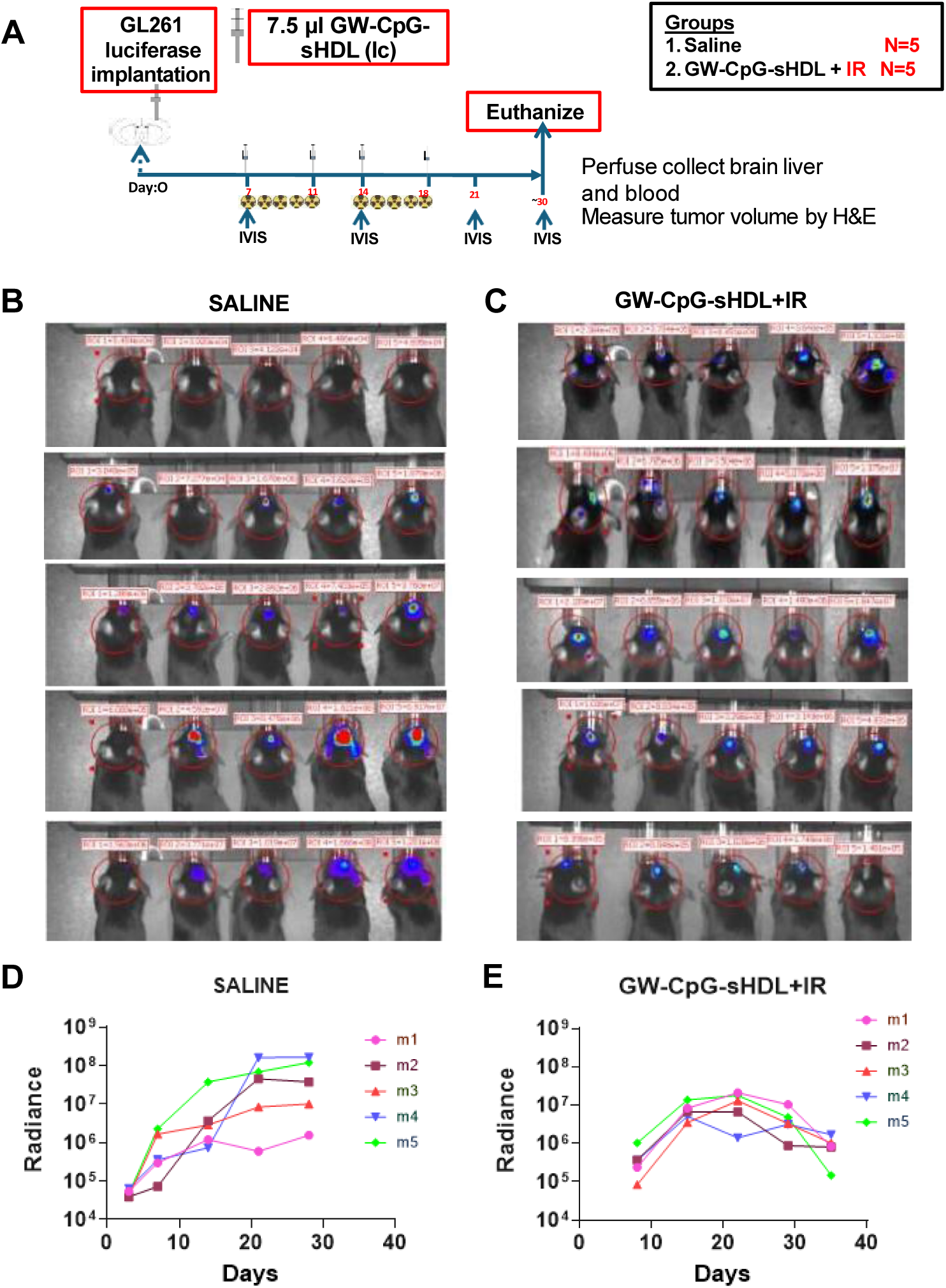
Tumor progression and the efficacy of GW-CpG-sHDL + IR treatment. A. Schematic representation of the timed treatment timeline for saline and GW-CpG-sHDL + IR groups. Tumors were implanted on Day 0. Treatment with GW-CpG-sHDL and irradiation began on Day 7 and continued until Day 18, with irradiation (IR) administered at specified time points. B. Bioluminescence imaging (BLI) showing tumor progression in saline-treated group C. (BLI) showing tumor progression in GW-CpG-sHDL + IR-treated mice over the treatment period. Each row represents BLI data from days 3,7,14,21 and 28 for the saline group while for GW-CpG-sHDL treatment group days 7,14,21 and 28 and 32, highlighting tumor radiance for all mice in both groups. D-E. Individual bioluminescence radiance plots for mice in the saline group (left panel) and the GW-CpG-sHDL + IR group (right panel). Radiance (photons/sec) was plotted over time, showing progressive tumor growth in the saline group and tumor suppression in the GW-CpG-sHDL + IR group.

### Tumor volume

The results provide a comparison of tumor characteristics between saline-treated and GW-CpG-sHDL-treated groups, highlighting significant differences in tumor volume, radius, and cross-sectional area. In the saline group, tumor volumes were found to have an average volume of 23.65 mm (Table 1). Individual measurements for saline-treated mice (m1, m2, m3) exhibited volumes ranging from 17.39 mm³ to 27.43 mm³, demonstrating larger tumor sizes.

**Table 1.** Tumor metrics between saline and GW-CpG-sHDL treated groups.

|  | Area (mm<sup>2</sup>) | Radius (mm) | Volume (mm<sup>3</sup>) |
| --- | --- | --- | --- |
| **Saline m1** | 20.57 | 2.56 | 27.43 |
| **Saline m2** | 13.04 | 2.04 | 17.39 |
| **Saline m3** | 19.61 | 2.5 | 26.14 |
| **Mean ± SEM** | 17.74 ± 2.37 | 2.37 ± 0.16 | 23.65 ± 3.15 |
| **Treated m1** | 1.87 | 0.77 | 2.49 |
| **Treated m2** | 0.2 | 0.25 | 0.26 |
| **Treated m3** | 3.71 | 1.09 | 4.94 |
| **Mean ± SEM** | 1.93 ± 1.01 | 0.70 ± 0.24 | 2.56 ± 1.35 |

Conversely, GW-CpG-sHDL treated groups showed markedly reduced tumor metrics, with an average volume of 2.56 mm³, indicating the reduction in tumor sizes with the treatment. Individual measurements for treated mice revealed volumes as low as 0.26 mm3. These results indicate an effective treatment strategy for glioma, with substantial reductions in overall tumor size compared to the saline group.

## Discussion

Glioblastoma remains an extremely difficult disease to treat to date. Despite radiation therapy, chemotherapy, and surgical resection, these tumors always recur. Glioblastoma patients have an average survival of 14 months post-diagnosis even with advanced treatments. The highly invasive nature of the tumor makes complete resection of the tumor unrealistic, and the development of resistance to temozolomide highlights the need for new treatment strategies to be developed for GBM. Considering the high recurrence rate, therapies capable of driving immunological memory against the tumor are particularly attractive for GBM.

Overcoming drug delivery challenges to the central nervous system, particularly for glioblastoma, is a major focus in medical research. The blood-brain barrier (BBB) restricts therapeutic agents, limiting treatments like surgery, radiotherapy, and chemotherapy. Researchers are exploring nanoparticle-based systems to address these limitations and reduce systemic toxicities.^[35–37]^ For example, Apolipoprotein E (ApoE) is being studied to enhance drug delivery to the brain.^[36]^ ApoE interacts with BBB receptors, which are also present on glioblastoma cells. ApoE-functionalized nanosystems, like polymersomes, can navigate the BBB and target glioblastoma, improving therapeutic delivery. Additionally, apoptotic proteins like granzyme B (GrB) show potential for glioma immunotherapy.^[36]^ GrB, from natural killer cells and cytotoxic T lymphocytes, induces cell death through mitochondrial attack. Recent studies use ApoE-directed, redox-responsive virus-mimicking polymersomes to deliver GrB to glioblastoma cells, enhancing delivery and opening new possibilities for targeted immunotherapy.^[36]^

In this study, we developed a synthetic high-density lipoprotein nanodisc platform aimed at reducing cholesterol concentration within the tumor microenvironment and delivering CpG adjuvants to drive a robust antitumor immune response. One of the central components of this platform is the Liver X Receptor agonist incorporated into the sHDL membrane. LXR agonists initially received significant research interest for their applications in atherosclerosis and cardiovascular disease^[16]^. Still, a growing body of literature demonstrates the utility of LXR agonists in various cancers.^[12, 14, 15, 18, 38–40]^ We chose to investigate 3 different LXR agonists in our experiments, namely T0901317, RGX-104, and GW3965. The antitumor effects of LXR agonists stem from several avenues, including depleting cells of cholesterol, upregulation of proteins involved in cell cycle arrest, and reversing the immunosuppressive effects of myeloid-derived suppressor cells (MDSCs) in the tumor microenvironment. Using sHDL nanodiscs as a delivery platform was particularly interesting given the established synergistic effect between LXR agonists and sHDL, wherein sHDL acts as an acceptor of excess cholesterol following upregulation of the ABCA1 transporter. By decorating the nanodiscs with CpG oligonucleotides, established TLR9 agonists, we aimed to stimulate the immune system further and enable immunological memory against glioma in the case of recurrence.

Our results highlight the successful synthesis and characterization of sHDL loaded with the different LXR agonists and cholesterol-modified CpG oligonucleotides. While each formulation was similar in size, morphology, and incorporation efficiency of the LXR agonist, there were important differences in the stability of the LXR agonist in the sHDL membrane. All 3 LXR agonists used in this study are hydrophobic compounds, with T0901317 having the highest aqueous solubility at ∼10µg/mL compared to ∼1µg/mL for RGX-104 and GW3965.^[18–20]^ It’s likely that RGX-104 and GW3965 were more stably associated with the sHDL membrane, given their greater hydrophobicity.

Subsequent testing of each formulation in glioma cells demonstrated effective delivery of LXR agonists and upregulation of one of the key downstream targets of LXR activation, ABCA1. This resulted in a ∼2-fold increase in the amount of cholesterol removed from cells compared to sHDL alone and a dose-dependent cytotoxic effect. Our group has previously employed sHDL nanodiscs as a delivery vehicle for LXR agonists for the treatment of atherosclerosis and diabetic nephropathy.^[21–23]^ The synergistic effect between sHDL and LXR agonists has been previously demonstrated in a number of macrophage and kidney cell lines, so our present findings in glioma cells are in agreement with existing literature and show that this strategy translates across a variety of cell types. Our studies on cell cycle analysis of both mouse and human glioblastoma cells following GW3965 and GW-CpG-sHDL treatment revealed an increase in the sub-diploidal population, representing cells with substantial DNA damage indicative of a late apoptotic stage relative to cycling cells. Furthermore, our findings confirm that GW-CpG-sHDL-induced cell death is mediated by caspase-3/7 activation in both mouse and human glioblastoma cells. Importantly, neither GW3965 nor GW-CpG-sHDL impacted cell cycle checkpoints, cell cycle phases, or caspase-3/7 activity in normal human astrocytes. Thus, GW-CpG-sHDL selectively increases the sub-diploidal population in glioblastoma cells, supporting a mechanism involving caspase-3/7 mediated apoptosis while sparing normal human astrocytes.

Extending our investigation to glioma-bearing mice, we observed statistically significant improvements in median survival for treatment groups compared to control saline-treated mice. In particular, 60% of mice receiving GW-CpG-sHDL nanodiscs in combination with radiation survived long-term (MS=not reached). This showed greater efficacy in comparison to either treatment administered individually. Rechallenging these mice in the contralateral hemisphere resulted in all mice surviving at least 45 days, with 2 out of 3 mice showing no signs of tumor development 60 days post-rechallenge. The significant improvement in survival observed when combining these strategies is likely due to targeting multiple pathways to induce cell death (DNA damage, cholesterol depletion, immune stimulation). The results from control and treated mice at day 30 post-tumor implantation;12Days after the last treatment demonstrate that treatment significantly reduces tumor burden while preserving normal brain tissue integrity. Enhanced CD8+ and CD4+ T-cell infiltration in treated tumors suggests a robust adaptive immune response. Lower Iba1 staining indicates decreased microglial activity, suggesting a shift in the tumor microenvironment toward an anti-tumor state. These findings underscore the multifaceted impact of the treatment in modulating the tumor microenvironment and promoting tumor regression (Table 1). Prior work has demonstrated that radiation therapy can increase the number of MDSCs present in the tumor microenvironment by augmenting the levels of CCL2, CCL5, and CSF-1.^[38]^ More MDSCs lead to higher levels of nitric oxide, TGF-β, Arg1, and IL-10, ultimately suppressing the activity of CD4+ and CD8+ T-cells. Several groups have demonstrated the immunosuppressive effects of MDSCs are reversible upon treatment with LXR agonists such as GW3965 and RGX-104.^[14, 24, 38, 39, 41]^ Tavazoie et al. demonstrated that LXR-mediated increases in the expression of ApoE lead to apoptosis of MDSCs and improved tumor immunity.^[14]^ Our group has previously used sHDL-CpG nanodiscs to deliver neoantigen peptides or docetaxel for treating GBM. In both studies, significant increases in CD8+ T-cells were observed in GL261 tumors.^[24, 25]^

Despite the benefits in survival observed with GW-CpG-sHDL nanodiscs combined with IR, there are challenges to be addressed with this therapeutic strategy. While several clinical studies demonstrate the safety of synthetic high-density lipoproteins, repeated use of early generation LXR agonists (such as T0901317 and GW3965) have been linked to liver toxicity due to increased triglyceride synthesis. In the present study, delivery of LXR agonists with sHDL nanodisc did not result in any changes in liver pathology or levels of enzymes indicative of healthy liver function. Similar observations have been reported from our group in which sHDL delivery of LXR agonists is well tolerated and mitigates increases in triglyceride levels.^[21–23]^ The assessment of the functional recovery of mice post-therapy conducting behavioral studies, falls beyond the scope of the current study. One of the key challenges in using GW-CpG-sHDL nanodiscs for GBM treatment is the requirement to bypass the blood-brain barrier in order to achieve therapeutic concentrations in the tumor. Intracranial dosing poses a significant barrier, and developing a nanodisc formulation that can be administered systemically while still reaching the tumor would be of much greater benefit. A number of different strategies for transporting nanocarriers across the BBB, such as conjugation to ApoE-mimetic peptides, RVG peptides, and anti-transferrin receptor antibodies, have been investigated by other groups,^[42–44]^ and these may offer an opportunity to improve upon this therapeutic strategy for GBM in future work.

## 3. Conclusions

In summary, we have reported a new adjuvant treatment strategy for GBM consisting of synthetic high-density lipoprotein nanodiscs delivering a Liver X Receptor ^[45]^cholesterol removal from glioma cells as a result of upregulation of the ABCA1 transporter and subsequent loading of cholesterol to sHDL nanodiscs. Treatment of glioma-bearing mice with GW-CpG-sHDL nanodiscs resulted in modest improvements in median survival. When combined with standard-of-care radiation therapy, nanodisc treatment greatly improved survival, with 60% of mice surviving long-term and developing immunological memory against the tumors when rechallenged. Overall, these findings highlight the value of targeting cholesterol metabolism and reversing immunosuppression in the development of new cancer therapies.

## 4. Experimental Section

### Reagents

The LXR agonists, T0901317 (T0), RGX104 (RGX), and GW3965 (GW3), were purchased from Cayman Chemical (Ann Arbor, MI). The ApoA-I mimetic peptide, 22A (PVLDLFRELLNELLEALKQKLK), was synthesized by Genscript Inc. (Piscataway, NJ). 1,2-dimyristoyl,sn-glycero-3-phosphocholine (DMPC) was purchased from Avanti Polar Lipids (Alabaster, AL). Cholesterol-CpG 1826 was synthesized by Integrated DNA Technologies Inc. (Coralville, IA). Cholesterol [1,2-^3^H(N)] was purchased from American Radiolabeled Chemicals (Saint Louis, MO). Anti-CD68 Abcam (ab125212), Anti-MBP (a.a. 82-87) Millipore (MAB386), Anti-GFAP Millipore (AB5541), Anti-Nestin NB1001604, Anti cd80 (Novas, NBP2-25255SS, 1:500), Anti TREM119 (Abcam,ab209064, 0.1 - 0.5 µg/ml), Anti-CD8a (230-247) (HistoSure, (HS-361 003), 1:100), Anti-Iba1 (Abcam, ab178846, 1:2000. Additional reagents used were of analytical grade and obtained from commercial suppliers.

### Preparation of LXR-CpG-sHDL Nanodiscs

LXR agonist, DMPC, and 22A were dissolved in acetic acid at a ratio of 1.5:30:15 (by mass in mg). This solution was then frozen in liquid nitrogen and freeze-dried for 24 hours. The resulting powder was rehydrated in 1 mL of pH 7.4 PBS and subjected to 3 cycles of heating at 50°C and cooling on ice (5 minutes each). The pH of each formulation was adjusted to 7.4 using NaOH. Unincorporated LXR agonist was removed by passing the formulation through a 7k MWCO Zeba desalting column. The chol-CpG was incorporated into the purified LXR-sHDL by incubating 34uL of a 2.24mg/mL stock solution with 1 mL of LXR-sHDL (final CpG concentration of 75 ug/mL) for 1 hour at room temperature on an orbital shaker at 100 rpm. The final solution was filtered using a 0.22µm syringe filter prior to use.

### Characterization of LXR-CpG-sHDL Nanodiscs

The incorporation efficiency of LXR agonists in sHDL nanodiscs was determined using LC-MS. Formulations before and after the removal of free drug were diluted 500-fold in methanol, and 5uL was injected on an Acquity UPLC H Class equipped with a QDa detector (Waters, Milford, MA). Samples were separated on an Acquity UPLC BEH C18 column (1.7 µm, 2.1×50 mm) using a gradient elution starting at 50:33:17 (A:B: C) and shifting to 0:67:33 (A:B: C) by 2.5 minutes, holding for 2 minutes, and returning to the starting composition at 5 minutes. The mobile phases A, B, and C were delivered at a flow rate of 0.3 mL/min and consisted of 0.1% formic acid in water, acetonitrile, or methanol, respectively. The LXR agonists were detected at the following M/Z ratios in negative mode with a cone voltage of 15V: GW3965 (580.09), RGX104 (594.25), T0901317 (480.18). The incorporation efficiency was calculated as the concentration of LXR agonist after purification divided by the concentration prior to purification.

Incorporation of chol-CpG into the nanodiscs was verified using size exclusion chromatography on a Waters 1525 HPLC (Waters, Milford, MA) with a Tosoh TSK gel G3000SWXL 7.8 mm × 30 cm column (Tosoh Bioscience, King of Prussia, PA) with PBS as the mobile phase delivered at 1 mL/min.

### Particle Size

The size of the sHDL nanodiscs was determined via dynamic light scattering (DLS) on a Malvern Zetasizer after diluting each formulation to 1 mg/mL (22A peptide concentration) with PBS.

### Drug Release

Each formulation was diluted to an LXR agonist concentration of 0.1 mg/mL in PBS containing BioBeads adsorbent media (Bio-Rad) to create sink conditions. Formulations were incubated at 37°C with shaking at 100 rpms, and aliquots were collected over the course of 48 hours. Aliquots were diluted with methanol, and the amount of drug remaining after each time point was quantified using LC-MS.

### Electron Microscopy

The morphology of each formulation was observed using transmission electron microscopy (TEM) with negative staining. sHDL samples were diluted to 10µg/mL (22A peptide concentration), and 4 uL was adsorbed for 1 minute to a glow-discharged 400-mesh copper grid with formvar carbon film (Electron Microscopy Sciences). The grids were washed and negative stained with 0.1% uranyl acetate. Samples were imaged on an FEI Morgagni electron microscope run at 100 kV at a magnification of 22,000X.

### Cell Lines and Culture Conditions

The GL261 cells were obtained from Creative BioArray (CSC-C9184W). The GL26 murine glioma cells were obtained from National Cancer Institute (NCI; Bethesda, MD). Cells derived from GL26 cells i.e. GL26-EGFRvIII, modified to express shp53, NRAS and EGFRvIII, genetic modifications relevant for human glioblastoma.^[46]^ Both GL26 EGFRvIII and GL261 cells were grown in Dulbecco’s Modified Eagle Medium (DMEM) (Gibco, 12430-062) supplemented with 10% fetal bovine serum (FBS) (Gibco, 10437-028) and 1X antibiotic-antimycotic (Gibco, 15240-062) solution. Cells were maintained in a humidified incubator at 37°C and 5% CO_2_.

### Primary Human GBM Cells and Culture Conditions

HF2303 primary human GBM cancer stem-cells were provided by Dr. Tom Mikkelsen, M.D. (Department of Neurology, Henry Ford Hospital, Detroit, MI)^[47]^ MGG8 glioma cells (original male donor) were the kind gift of Dr. Samuel Rabkin, Department of Pathology, Harvard Medical School.^[45]^ Both HF2303 and MGG8 cells were grown in Dulbecco’s modified Eagle medium (DMEM) (Gibco, 12430-062) supplemented with 10% fetal bovine serum (Gibco,10437-028), 1X L-glutamine (100X; Gibco,25030-081), 1X MEM Non-Essential Amino Acids (Gibco, 11140-050), 1X Sodium pyruvate (Gibco, 11360-070) and antibiotic-antimycotic (1×) (Gibco, 15240-062) solutions. Cells were maintained in a humidified incubator with 5% CO_2_ at 37^0^C and passaged every 2-3 days.

### RT-qPCR

GL261 cells were plated in 6-well plates at 600,000 cells/well density. Cells were treated with DMSO, sHDL, free LXR agonist, or LXR agonist loaded into sHDL for 24 hours at an LXR agonist concentration of 5 µM. RNA was isolated from cells using a Qiagen RNeasy isolation kit. Purified RNA was then reverse transcribed using a High-Capacity RNA-to-cDNA Kit (Applied Biosystems), and qPCR was performed using TaqMan Gene Expression Assays (Applied Biosystems). The relative quantities of LXRα, LXRβ, and ABCA1 were determined using the ΔΔCt method with 18S as the housekeeping gene.

### Western Blotting

GL261 cells were plated in 6-well plates at a density of 600,000 cells/well. Cells were treated with DMSO, sHDL, free LXR agonist, or LXR agonist loaded into sHDL for 24 hours at an LXR agonist concentration of 5 µM. After treatment, the cells were lysed with RIPA lysis buffer (50 mM Tris-HCl, 150 mM NaCl, 1% NP-40, 0.5% sodium deoxycholate, 0.1% SDS (Boston BioProducts) supplemented with protease and phosphatase inhibitor cocktails (Thermo Scientific). The protein concentration of each sample was determined by BCA assay (Pierce), and equal amounts of protein extracts were separated on 4–12% NuPAGE Bis-Tris Mini Gel (Invitrogen) and then transferred to a nitrocellulose membrane (GE Healthcare) with XCell II Blot Module (Invitrogen). The membrane was probed with antibodies for LXRα, LXRβ, ABCA1, and Actin, followed by secondary antibodies conjugated to horseradish peroxidase. The protein bands were visualized using a Fluorchem M Imaging system (ProteinSimple).

### Cholesterol Efflux

GL261 cells were plated in a 24-well plate at a density of 100,000 cells/well in media containing 1 µCi/mL of cholesterol [1,2-^3^H(N)]. After 24 hours, the cells were washed twice with PBS and treated with DMSO, sHDL, 1 µM free LXR agonist, or 1 µM LXR agonist loaded into sHDL in DMEM containing 10% lipoprotein deficient serum. After treating the cells for 24 hours, media and cell lysates were collected and diluted with a scintillation cocktail (200µL media/cell lysate + 1mL scintillation cocktail). The radioactivity of each sample was measured on a liquid scintillation counter (Perkin Elmer, Waltham, MA). The %cholesterol efflux in each group was calculated by dividing the media counts by the sum of the media and cell lysate counts. The %cholesterol efflux from the DMSO-treated cells (negative control) was subtracted from each group.

### Cell Survival Assay

GL261 cells were plated in a 96-well plate at a density of 3000 cells/well overnight. The media was removed and replaced with fresh media containing DMSO, sHDL, free LXR agonist (at 1, 5, or 10 µM), or LXR agonist loaded into sHDL (at 1, 5, or 10 µM). After 24 hours, cell viability was determined via a CellTiter96 Aqueous One Solution (Promega, Madison, WI) according to the manufacturer’s instructions. Cell viability was normalized to DMSO-treated cells. In another set of study, normal human astrocytes, mouse glioblastoma GL26 EGFRvIII along with human patients-derived glioblastoma cells HF2303 and MGG8 cells were plated at a density of 1000 cells per well in a 96-well plate (Fisher, 12-566-00) 24h before treatment. Cells were then incubated with either saline, empty-sHDL, CpG-HDL, free-GW3965 (LXR agonist; SelleckChem, S2630) and [GW3965 + CpG] loaded-sHDL (GW-CpG-HDL) alone or in combination with radiation at 1, 2.5 or 5 µM doses for 72 hours in triplicate wells per condition. sHDLs without GW3965 were tested in the same peptide and lipid concentrations as CpG-sHDL-GW with specified GW3965 concentration. All the mouse and patient-derived glioma cells were pretreated with 2h prior to irradiation with 3 Gy for mouse and 14 Gy of radiation for human cells, respectively. Cell viability was determined with CellTiter-Glo 2.0 Luminescent Cell Viability Assay (Promega, G9242) following manufacturer’s protocol. Resulting luminescence was read with the Enspire Multimodal Plate Reader (PerkinElmer). Cell viability was normalized to solvent-treated controls. Data were represented graphically using the GraphPad Prism 9.0 software, and statistical significance was determined by one-way ANOVA followed by Tukey’s test for multiple comparisons.

### Caspase 3/7 activation assay

The caspase-3/7 activation assay was performed following the manufacturer’s protocol for the Caspase–Glo® 3/7 Apoptosis Assay System (G8091; Promega, Madison, WI, USA). Briefly, normal huma astrocytes, mouse and human glioblastoma cells were seeded in a 96-well plate at a density of 2,000 cells/well. Twenty-four hours after seeding, the cells were treated with phosphate-buffered saline (DPBS), empty nanoparticles (CpG-sHDL), free LXR agonist (GW3965), or GW3965-loaded nanoparticles (GW-CpG-sHDL). Staurosporine (0.5µM) and DMSO (20%) were used as positive and negative controls, respectively, for 12 hours. After 12 hours of incubation, the plates were centrifuged at 2,000 × g for 5 minutes, and 100 µl of the supernatants were transferred to a clean, opaque, flat-bottom 96-well plate. Reconstituted Caspase–Glo® 3/7 reagent was added to each well, and the plate was incubated at room temperature with shaking for 30 minutes. Luminescence was measured using a Tecan SPARK multimode microplate reader (Tecan Austria GmbH, Grödig, Austria). The data were normalized to protein concentration and presented as bar diagrams.

### Cell cycle analysis

For cell cycle analysis, both human and mouse glioblastoma cells were seeded in T12.5 cm² tissue culture flasks at a density of 1 × 10⁵ cells/flask. Twenty-four hours after seeding, the cells were treated with phosphate-buffered saline (DPBS), empty nanoparticles (CpG-sHDL), free LXR agonist (GW3965), or GW3965-loaded nanoparticles (GW-CpG-sHDL). Staurosporine (0.5 µM) was used as a positive control for 12 hours. Following 48 and 72 hours of incubation, the treated cells were fixed in 70% (v/v) ice-cold ethanol and kept overnight at 4 °C. After washing twice with PBS, the cells were incubated with RNase (250 µg/mL) at 37 °C for 60 minutes (Sigma Chemical Company, St. Louis, MO). The cells were then stained with propidium iodide (PI, 20 µg/mL). The cell cycle distribution and the percentage of apoptotic cells were analyzed using a ZE5 cell analyzer (Bio-Rad, California, USA), with 10,000 events collected per sample. Appropriate gating was applied to select the single-cell population, and the same gate was used across all samples to ensure standardized measurements.

### Tumor Implantation

C57BL/6 strain C57BL/6NTac was purchased from Taconic (cat# B6-F). All procedures involving mice were performed in accordance with policies set by the Association for Assessment and Accreditation of Laboratory Animal Care (AAALAC) and the Institutional Animal Care and Use Committee (IACUC) at the University of Michigan (PRO00011290). Mice were anesthetized with ketamine (75mg/kg, i.p) and dexmedetomidine (0.5mg/kg, i.p) before stereotactic implantation of GL261(50K) cells in the right striatum. The implantation coordinates were 1.0 mm anterior and 2.0 mm lateral from the bregma and 3.5 mm ventral from the dura, with an injection rate of 1 μL/min. After implantations mice were given combination of buprenorphine (0.1mg/kg, s.c) and carprofen (5mg/kg, s.c) for analgesia.^[48]^

One week post implantations mice were randomly divided into the following groups: Saline, Irradiation (IR), Liver X receptor agonist (LXR agonist, GW3965), LXR agonist (GW 3965)+IR, GW-CpG-sHDL, GW-CpG-sHDL + IR. Upon reaching the symptomatic tumor stage, blood from these mice was collected to perform the serum chemistry, and further mice were perfused with tyrodes solution to collect the brain, liver, and spleen.

### *In Vivo* Radiation

The groups treated with radiation alone or combined with LXR agonist GW3965 or GW-CpG-sHDL were subjected to an irradiation (IR) dose of 2 Gy for 5 days/week for 2 weeks for a total of 20 Gy of ionizing radiation. Irradiation treatment was done as follows: Mice were gently anesthetized with isoflurane and positioned under a copper orthovoltage source. The irradiation beam was targeted toward the brain while protecting the body with iron collimators. Irradiation treatment was performed at the University of Michigan Radiation Oncology Core.^[48, 49]^

### sHDL-CpG *In Vivo* Dosing

sHDL loaded with GW3965 and cholesterol-CpG were filtered through a 0.22 µM syringe filter and diluted to a GW3965 concentration of 1.5 mg/mL in sterile PBS. For the intracranial (i.c) dose, 7.5ul GW-CpG-sHDL was used. For the intraperitoneal injection (i.p) of the LXR agonist GW3965, 0.5mg/kg of the compound was used in 100 μL of PBS.

### Tumor Volume calculation

To calculate the tumor volume, the cross-sectional area of the tumor was measured using ImageJ software^[50]^. The histological image was loaded, and the scale bar was calibrated to mm. The tumor boundary was manually traced using the freehand tool, and the area was calculated in mm² using the Analyze > Measure function(^[50]^). The method assumes the tumor has a spherical geometry for volume approximation, ImageJ was used for area measurement, while standard mathematical formulas were applied for the volume approximation.

Assuming a spherical tumor shape, the volume was approximated using the formula:

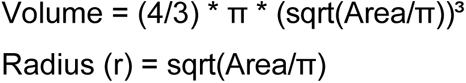

### Complete Blood Serum Biochemistry

Blood from tumor-bearing mice was taken from the submandibular vein and transferred to serum separation tubes (Biotang). Samples in the serum separation tubes were left at room temperature for 60mins to allow blood coagulation before centrifugation at 2000 rpm (400 x g). Complete serum chemistry for each sample was determined by in vivo animal core at the University of Michigan.

### Immunohistochemistry

For neuropathological assessment, brains fixed in 4% paraformaldehyde (PFA) were embedded in paraffin and sectioned 5µm thick using a microtome system (Leica RM2165). Immunohistochemistry was performed on brain sections by permeabilizing them with TBS-0.5% Triton-X (TBS-Tx) for 20 min. This was followed by antigen retrieval at 96 °C with 10 mM sodium citrate (pH 6) for 20 min. Then, the sections were allowed to cool at room temperature (RT), followed by washing 5 times with TBS-Tx (5 min per wash), and blocked with 5% goat serum in TBS-Tx for 1 h at RT. Brain sections were incubated with primary antibody GFAP (Millipore Sigma, AB5541, 1:200), MBP (Millipore Sigma, MAB386, 1:200), CD68 (Abcam, ab125212, 1:1000) or Nestin (Novus Biologicals, NB100-1604, 1:1000), Anti-Iba1(Abcam,(ab178846), 1:2000), CD8a (HistoSure, HS-361 003, 1:100), and CD4 (Cell Signaling, 48274S, 1:100) diluted in 1% goat serum TBS-Tx overnight at RT. On the following day, sections were washed with TBS-Tx 5 times. Brain sections labeled with GFAP, CD68, MBP, and Nestin were incubated with biotinylated secondary antibody diluted 1:1000 in 1% goat serum TBS-Tx in the dark for 4h. Biotin-labeled sections were subjected to 3, 3′-diaminobenzidine (DAB) (Biocare Medical) with nickel sulfate precipitation. The reaction was quenched with 10% sodium azide; sections were washed 3 times in 0.1 M sodium acetate, followed by dehydration in xylene, and coverslipped with DePeX Mounting Medium (Electron Microscopy Sciences). Images were obtained using brightfield microscopy (Olympus BX53) at 10X and 40X magnification.

Brain sections measuring 5µm in thickness, embedded in paraffin, were prepared from each experimental group to quantify tumor size. These sections were subsequently stained with hematoxylin and eosin (H&E).^[51]^ Specifically, sections containing tumor areas or derived from rechallenged brains (approximately 10-12 sections per mouse) were captured using brightfield microscopy (Olympus BX53) settings to acquire the images.

For histological assessment, livers were embedded in paraffin, sectioned 5µm thick using the microtome system, and H&E stained. Brightfield images were obtained using the Olympus MA BX53 microscope.^[51]^

### Statistical Analysis

For *in vitro* experiments, treatments were run in triplicates and compared using one-way ANOVA. For *in vivo* experiments, sample sizes were selected based on data from previous experiments done in our laboratories and published results from the literature. Animal experiments were performed after randomization. Kaplan-Meier survival curves were assessed using the log-rank (Mantel-Cox) test with Prism 8.1 (GraphPad Software). A p-value less than 0.05 was considered statistically significant.

## Supporting information

Supplementary Information is available as a PDF file

## 6. Acknowledgements

The laboratory of MGC is supported by the National Institutes of Health (NIH)/National Institute of Neurological Disorders and Stroke (NIH/NINDS) grants R37-NS094804, R01-NS105556, R01-NS122536, R01-NS124167, and R21-NS123879-01, and Rogel Cancer Center Scholar Award to M.G. Castro. This work has also been supported by The Department of Neurosurgery, The Pediatric Brain Tumor Foundation, Leah’s Happy Hearts Foundation, Ian’s Friends Foundation (IFF), Chad Tough Foundation, Smiles for Sophie Forever Foundation to M.G. Castro. TAH is funded by the Predoctoral Fellowship in Drug Discovery from the PhRMA Foundation. MY is supported by American Heart Association Postdoctoral Fellowship (24POST1196020). We would also like to acknowledge the University of Michigan In-Vivo Animal Core (IVAC) facility.

