## Supplementary Information is available as a PDF file for "HDL Nanodiscs Loaded with Liver X Receptor Agonist Decreases Tumor Burden and Mediates Long-term Survival in Mouse Glioma Model"

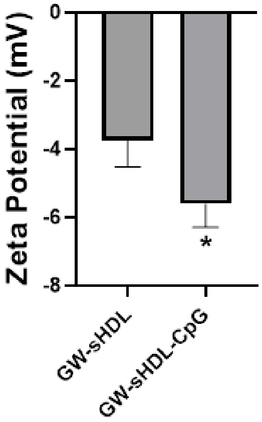


**Figure S1.** Zeta potential of GW-sHDL and GW-CpG-sHDL measured by DLS. (Mean ± SD, n = 3. *p < 0.05)

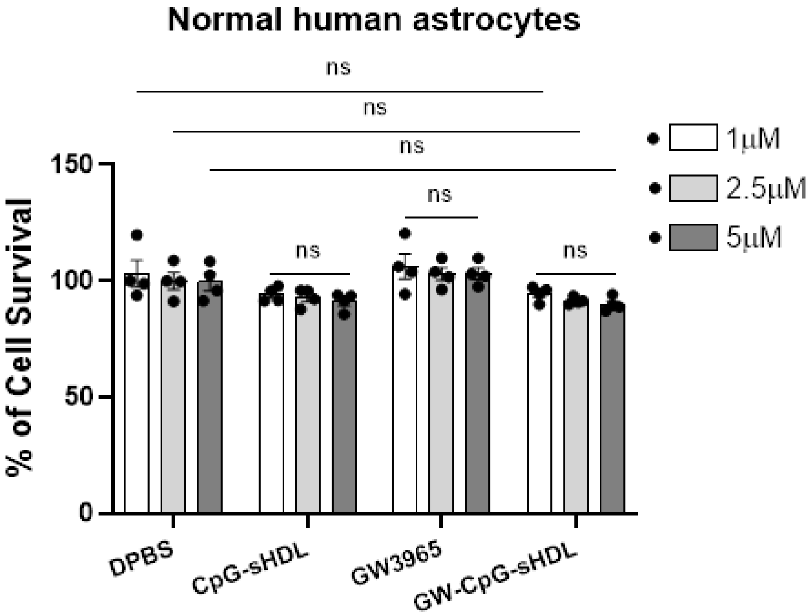


**Figure S2.** ***In vitro*** **cytotoxicity evaluation of LXR agonists (GW3965) and LXR agonist-loaded nanoparticles on normal human astrocytes.** Normal human astrocytes were treated with either free-GW3965, empty sHDL, CpG-sHDL, or GW-CpG-sHDL nanodiscs at concentrations of 1, 2.5 or 5 µM doses for 72 h. sHDLs without GW3965 were tested using the same peptide and lipid concentrations as GW-CpG-sHDL with corresponding GW3965 concentration. The bar plot shows the % of viable normal human astrocytes after treatment with each experimental condition. All statistical analysis was performed through unpaired t-test. Data represent Mean ± SEM (n = 3 biological replicates). Statistical notation: ns= not significant.

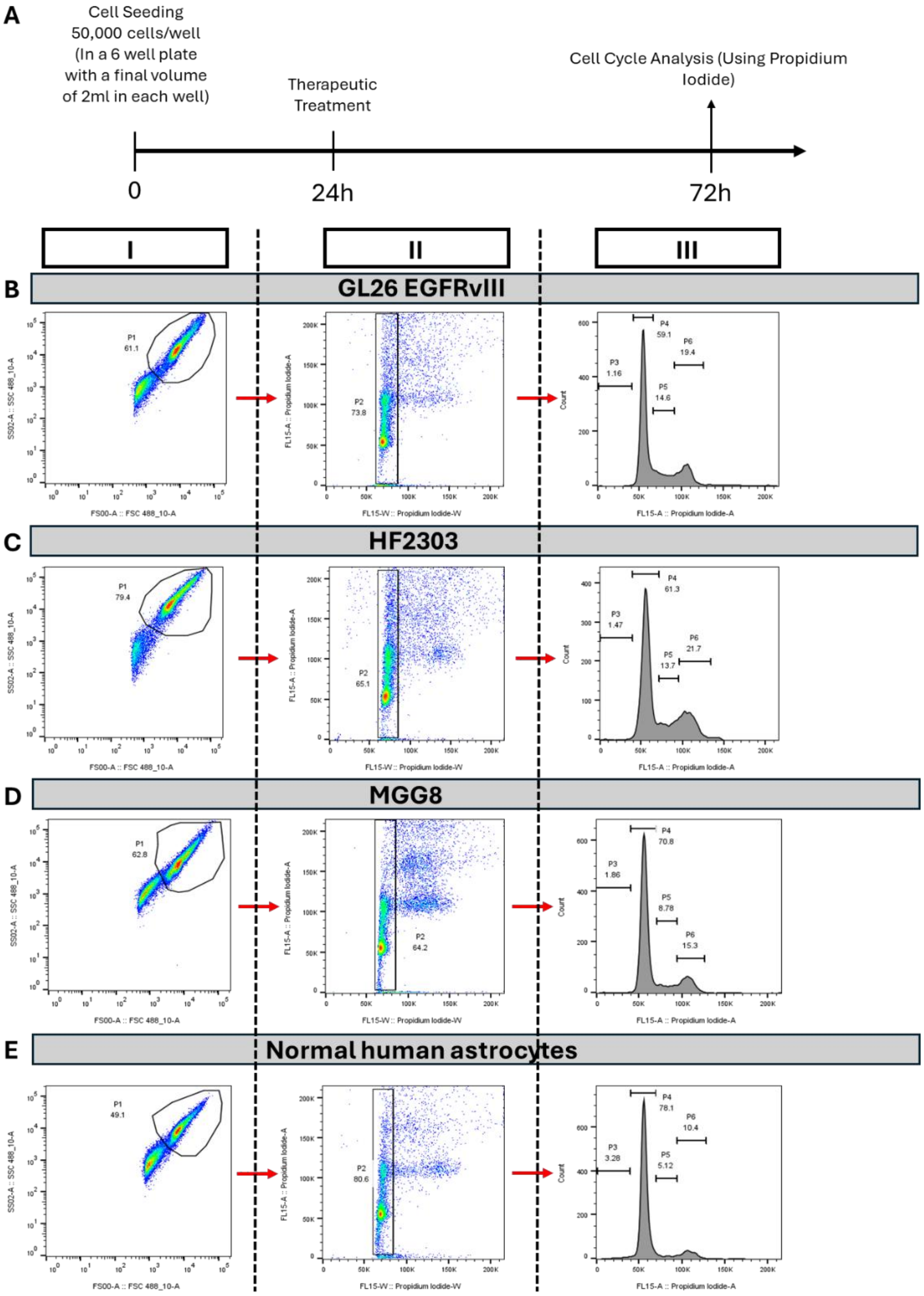


**Figure S3. Schematic illustration of experimental designs, timeline, and sequential gating strategies for flow cytometry analysis of cell cycle distribution. (A)** The schematic illustration of the timeline and experimental designs for the *in vitro* application of free-GW3965 and GW3965-CpG-HDL nanodiscs for cell cycle distribution assay on mouse glioblastoma (GL26 EGFRvIII), patient-derived glioblastoma cells (MGG8 and HF2303), and normal human astrocytes. Sequential gating strategy for flow cytometry analysis of cell cycle distribution for **(B)** GL26 EGFRvIII, **(C)** HF2303, **(D)** MGG8, and **(E)** normal human astrocytes are as follows: (I) propidium iodide-labeled cells were gated to exclude cellular debris. (II) doublet discrimination gating was performed to filter out cellular aggregates prior to analysis, and (III) cells were gated to determine the percentage of population in each cell cycle phase and presented as a histogram.

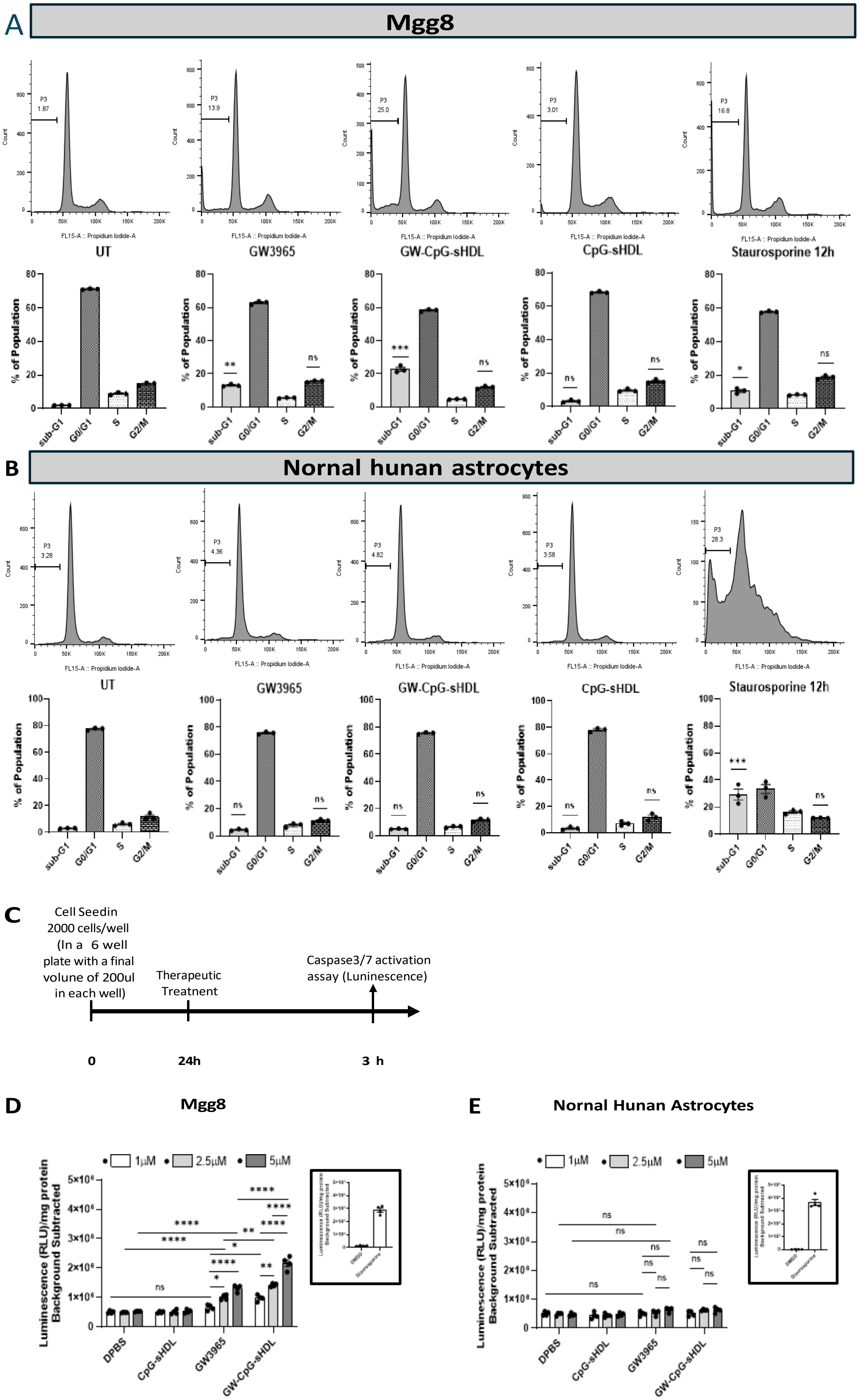

A
Mgg8
UT
GW3965
GW-CpG-sHDL
CpG-sHDL
Staurosporine 12h
FL15-A :: Propidium Iodide-A
Count
% of Population
sub-G1
G0/G1
S
G2/M
B
Nornal hunan astrocytes
C
Cell Seedin
2000 cells/well
(In a 6 well
plate with a final
volume of 200ul
in each well)
Therapeutic
Treatnent
Caspase3/7 activation
assay (Luninescence)
0
24h
3 h
D
Mgg8
E
Nornal Hunan Astrocytes
1μM
2.5μM
5μM
Luminescence (RLU)/mg protein
Background Subtracted
DPBS
CpG-sHDL
GW3965
GW-CpG-sHDL

**Figure S4. Cell cycle distribution and activation of caspase 3/7 in human glioblastoma and normal human astrocytes following treatment with LXR agonist and LXR agonist-loaded nanoparticles. (A)** MGG8 human glioblastoma and **(B)** normal human astrocytes cells were treated with either free-GW3965, empty sHDL, CpG-sHDL, or GW-CpG-sHDL nanodiscs at a concentration of 5 µM for 72 hours. Staurosporine (0.5 µM) served as a positive control for 12 hours. sHDLs without GW3965 were tested using the same peptide and lipid concentrations as those used in GW-CpG-sHDL treatments with corresponding GW3965 doses. Cells were harvested and fixed in 70% ethanol, stained with propidium iodide, and analyzed via flow cytometry. The percentage of cells in the sub-G1 phase (indicative of hypodiploid DNA content) is shown in each panel. Bar graphs display the “% of population” in different cell cycle phases, with statistical significance of other treatment groups compared to the untreated control. **(C)** Schematic illustration of the timeline for the *in vitro* application of free-GW3965 and GW-CpG-sHDL nanodiscs on mouse glioblastoma (GL26 EGFRvIII), patient-derived glioblastoma cells (MGG8 and HF2303), and normal human astrocytes. To assess caspase 3/7 activation, **(D)** MGG8 and **(E)** normal human astrocytes were treated with either free-GW3965, empty-sHDL, CpG-sHDL, or GW-CpG-sHDL nanodiscs at concentrations of 1, 2.5, or 5 µM for 12 hours. The Caspase-Glo 3/7 apoptosis assay showed a significant increase in normalized luminescence values in treated group compared to the untreated control, indicating pro-apoptotic bioactivity in a dose-dependent manner. Staurosporine (0.5 µM) and DMSO (20%) were used as positive and negative controls, respectively, for 12 hours (data presented in the inset). Values represent the Mean ± SEM of three independent experiments with four replicates per condition. Data were analyzed using one-way ANOVA followed by post hoc Tukey's test and unpaired Student’s t-test to assess significant differences between groups. Statistical significance relative to the untreated control is indicated as *$p < 0.05$, **$p < 0.01$, ***$p < 0.001$, ****$p < 0.0001$.

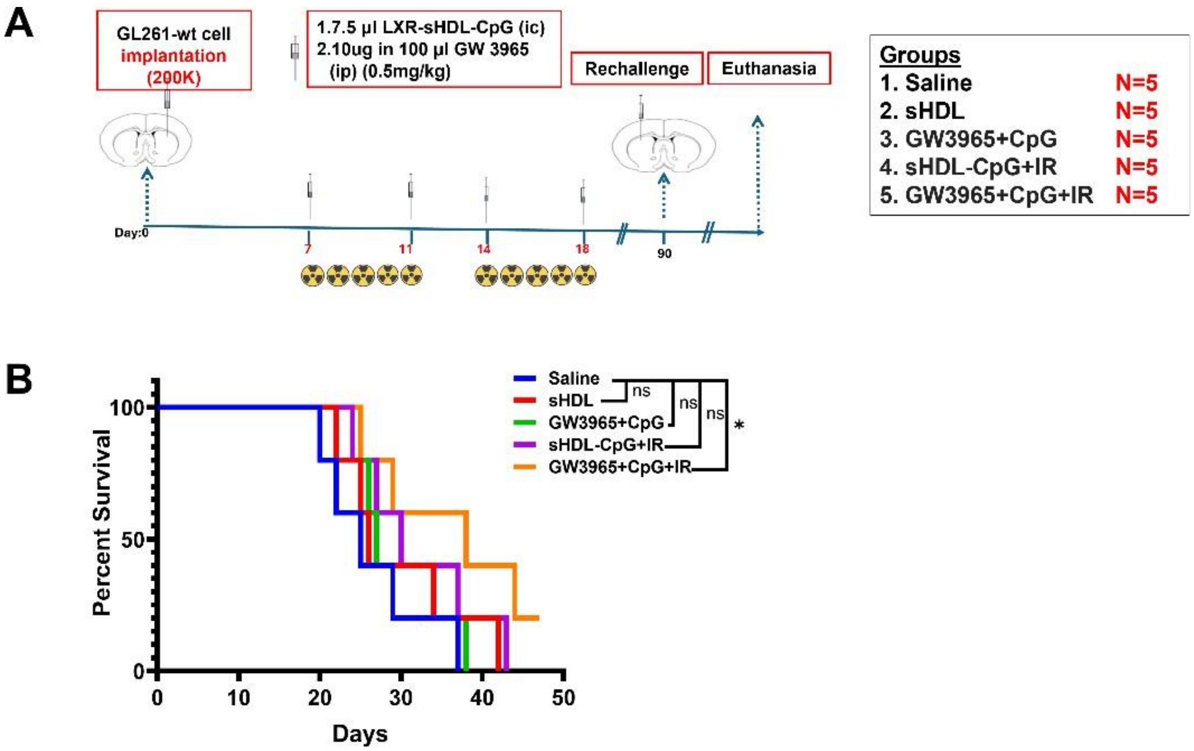


**Figure S5.** Synergistic effects of GW3865, CpG and IR. Outline of treatment schedule **(A)** with mice receiving saline, sHDL, free GW3965+CpG, sHDL-CpG + IR, or GW3965+CpG + IR at the indicated timepoints. Kaplan-Meier survival curves for each treatment group **(B).** n = 5 mice per group. *p < 0.05, **p < 0.01

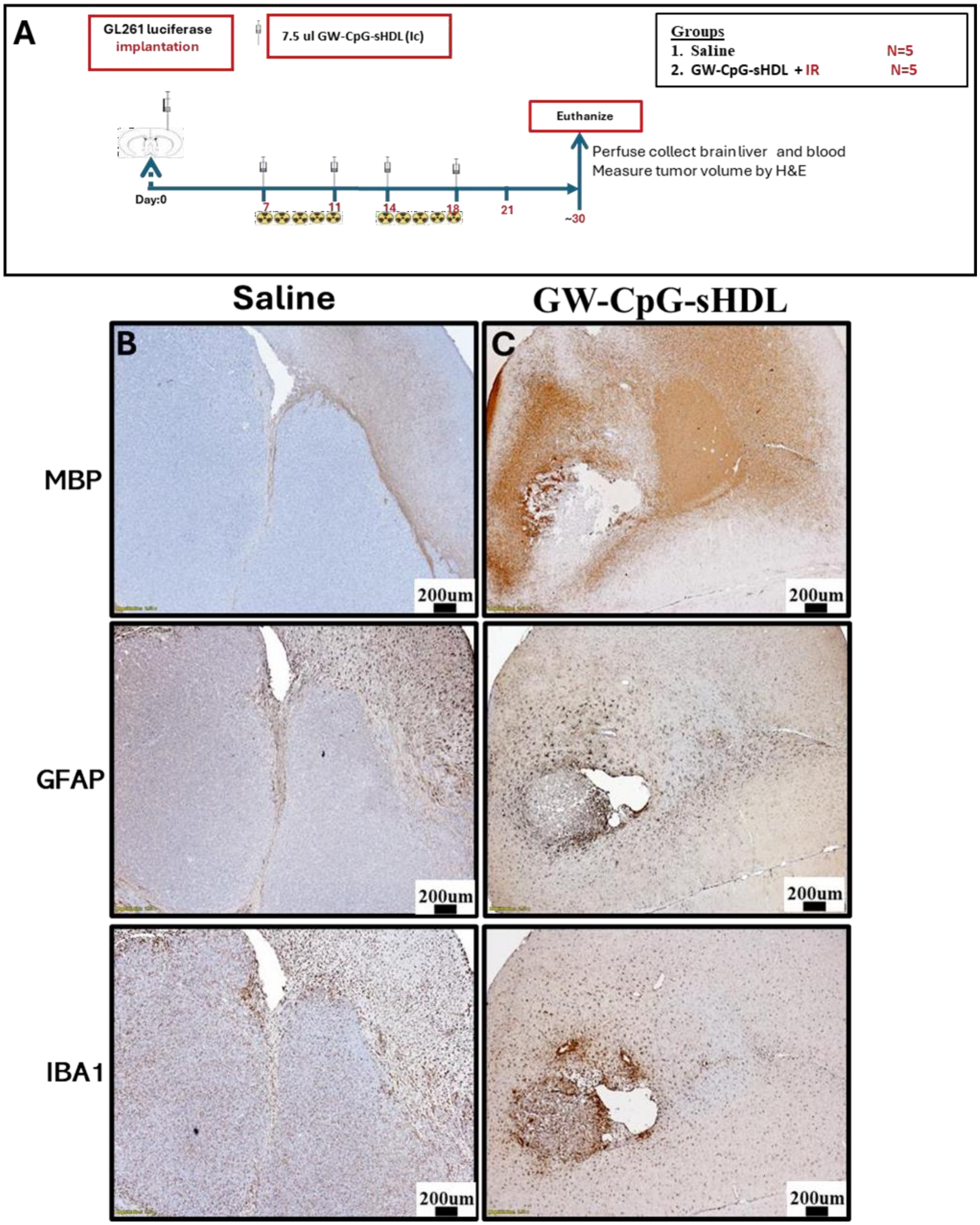


**Figure S6. Histological and immunohistochemical analysis of brain tumor sections from control and treated mice at day 30 post-tumor implantation;12Days after the last treatment. (A)** Outline of treatment schedule for the 30-day timed experiment. **(B and C)** MBP and GFAP staining indicate preserved myelin integrity and reduced astrocyte activation in treated tumors. Iba1 staining demonstrates reduced microglial activity in treated mice. Representative images from a single experiment consisting of independent biological replicates are displayed.

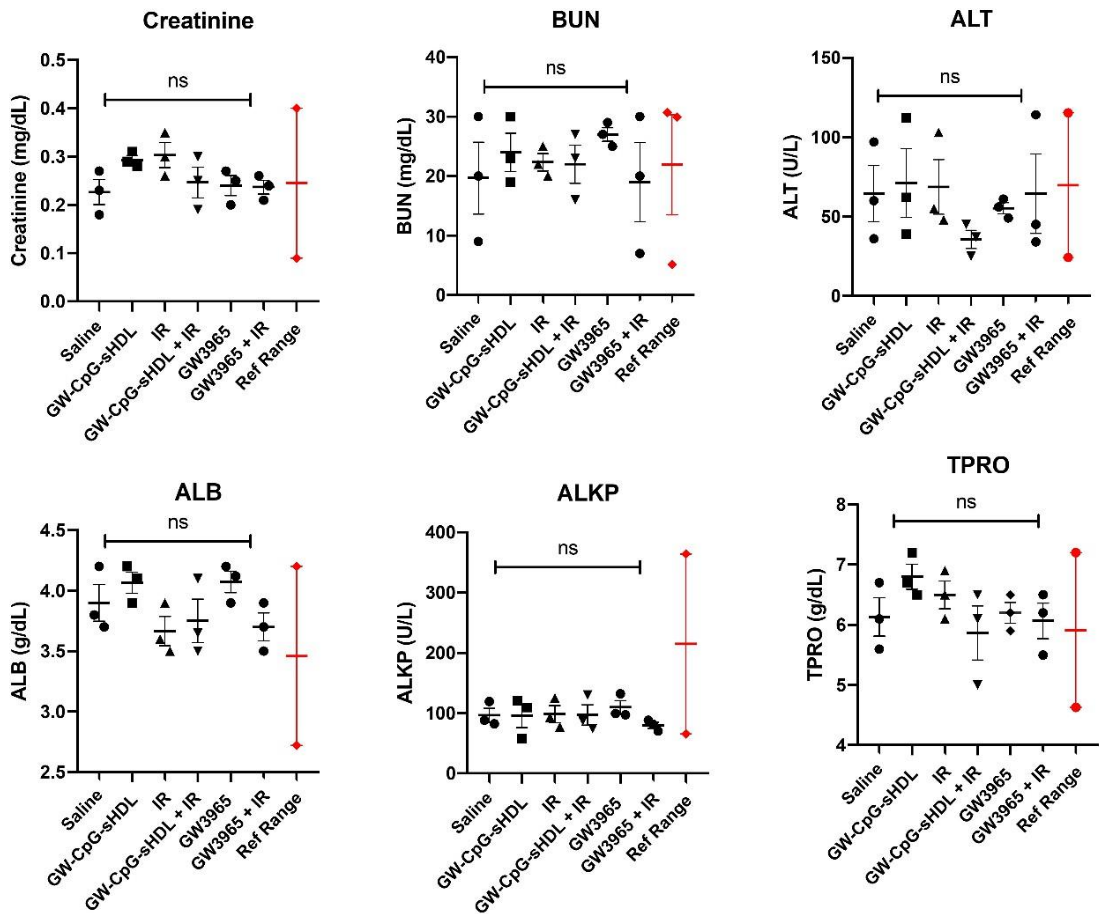


**Figure S7**. Mouse serum biochemical analysis following treatment in initial survival study. GL261 tumor-bearing mice from each treatment group exhibited normal serum biochemical parameters compared to saline-treated control. Serum was collected from tumor-bearing mice treated with saline, GW-CpG-sHDL, IR, or GW-CpG-sHDL + IR at 23 DPI. For each treatment group, levels of Creatinine, BUN, ALT, ALB, ALKP, and TPRO were quantified. The levels of different serum biochemical parameters between the treatment groups were compared and were found non-significant, $p > 0.05$ (n=3 biological replicates).

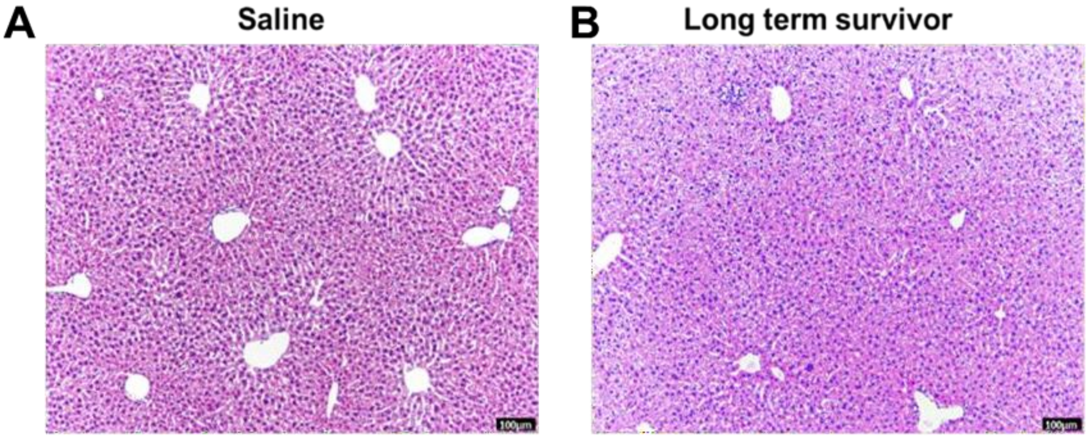


**Figure S8**. Histopathological assessment of livers from tumor-bearing mice treated with saline **(A)** or GW-CpG-sHDL + IR long term survivors which withstood the initial tumor challenge and survived over 3 months tumor-free **(B)**. H&E staining of 5 µm paraffin-embedded liver sections from saline and rechallenged long-term survivor groups was performed. No significant difference was observed in the histological features of hepatocytes and the stromal regions, both in the central and portal areas, when comparing the control saline group to the long-term survivor group. Representative images from a single experiment consisting of independent biological replicates are displayed. Black scale bars = 100 µm.
